# Understanding population structure in an evolutionary context: population-specific *F*_ST_ and pairwise *F*_ST_

**DOI:** 10.1101/2020.01.30.927186

**Authors:** Shuichi Kitada, Reiichiro Nakamichi, Hirohisa Kishino

**Author notes:** Corresponding author: Shuichi Kitada, Tokyo University of Marine Science and Technology, Minato-ku, Konan 4-5-7, Tokyo, 108-8477, Japan.

## Abstract

Populations are shaped by their history. It is crucial to interpret population structure in an evolutionary context. Pairwise *F*_ST_ measures population structure, whereas population-specific *F*_ST_ measures deviation from the ancestral population. To understand the current population structure and a population’s history of range expansion, we propose a representation method that overlays population-specific *F*_ST_ estimates on a sampling location map, and on an unrooted neighbor-joining tree and a multi-dimensional scaling plot inferred from a pairwise *F*_ST_ distance matrix. We examined the usefulness of our procedure using simulations that mimicked population colonization from an ancestral population and by analyzing published human, Atlantic cod, and wild poplar data. Our results demonstrated that population-specific *F*_ST_ values identify the source population and trace the evolutionary history of its derived populations. Conversely, pairwise *F*_ST_ values represent the current population structure. By integrating the results of both estimators, we obtained a new picture of the population structure that incorporates evolutionary history. The generalized least squares of genome-wide population-specific *F*_ST_ indicated that the wild poplar population expanded its distribution to the north, where daylight hours are long in summer, to seashores with abundant rainfall, and to the south with dry summers. Genomic data highlight the power of the bias-corrected moment estimators of *F*_ST_, whether global, pairwise, or population-specific, that provide unbiased estimates of *F*_ST_. All *F*_ST_ moment estimators described in this paper have reasonable process times and are useful in population genomics studies. The R codes for our method and simulations are available in the Supplemental Material.

---

Quantifying genetic relationships among populations is of substantial interest in population biology, ecology, and human genetics (Weir and Hill 2002). Appropriate estimates of population structure are the basis of our understanding of biology and biological applications, which vary from evolutionary and conservation studies to association mapping and forensic identification (Weir and Hill 2002). For such objectives, Wright’s *F*_ST_ (Wright 1951) is commonly used to quantify the genetic divergence of populations, and there have been many informative reviews of *F*_ST_ estimators (*e.g.*, Rousset 2004, 2007; Balloux and Lugon-Moulin 2002; Weir and Hill 2002; Beaumont 2005; Excoffier 2007; Holsinger and Weir 2009; Gaggiotti and Foll 2010; Bhatia *et al.* 2013). The traditional *F*_ST_ estimators have been defined as the ratio of the between-population variance to the total variance of allele frequencies (Wright 1965; Cockerham 1969, 1973; Weir and Cockerham 1984; Balloux and Lugon-Moulin 2002; Excoffier 2007; Holsinger and Weir 2009). An alternative approach for estimating population differentiation is to use population-specific *F*_ST_ estimators (Balding and Nichols 1995; Nicholson *et al.* 2002; Weir and Hill 2002; Weir *et al.* 2005; Gaggiotti and Foll 2010; Weir and Goudet 2017). Model-based Bayesian approaches, based on beta and/or Dirichlet distributions, for estimating population-specific *F*_ST_ have been proposed (Balding and Nichols 1995; Nicholson *et al.* 2002; Falush *et al.* 2003; Beaumont and Balding 2004). In addition to model-based methods, moment estimators of population-specific *F*_ST_ have been derived (Weir and Hill 2002; Weir and Goudet 2017). A large number of approaches exist for estimating *F*_ST_ that have different underlying assumptions (global, pairwise, or population-specific *F*_ST_) and the framework used, such as frequentist and/or Bayesian. There have been many comprehensive reviews of traditional and population-specific *F*_ST_ estimators, as indicated above, most of which were written from the viewpoint of theoretical issues. Conversely, there has been no formal comparative study to describe their differences in terms of the evolutionary scenarios that best describe the data. Although these issues are well understood among statistical geneticists and theoretical population geneticists, empirical researchers, particularly those working in non-model organisms, could benefit from studies that address the problem.

In practice, the *F*_ST_ value estimated from a set of population samples is called the global *F*_ST_ that measures population differentiation over all populations (*e.g.*, Pérez-Lezaun *et al.* 1997). Additionally, *F*_ST_ values between pairs of population samples (pairwise *F*_ST_, Reynolds *et al.* 1983) are routinely used to estimate population structure in human genetics (Pérez-Lezaun *et al.* 1997; Ramachandran *et al.*, 2005), conservation biology (Palsbøll *et al.* 2007; Schwartz *et al.* 2007), and evolutionary biology and ecology (*e.g.*, Hemmer-Hansen *et al.* 2013a; Therkildsen *et al.* 2013a; Geraldes *et al.* 2014; McKown *et al.* 2014a; Jorde *et al.* 2015; Rougemont *et al.* 2020). Divergent selection in an environmental gradient can influence population structure (Nosil *et al.* 2009; Orsini *et al.* 2013), and researchers have examined geographic distance and habitat differences between populations as explanatory variables that impact population structure estimated based on genome-wide (average over loci) pairwise *F*_ST_ values (*e.g.*, Rousset 1997; Bradbury and Bentzen 2007; Petrou *et al.* 2014; Jorde *et al.* 2015; Kitada *et al.* 2017). To identify the adaptive divergence of a gene among populations, locus-population-specific *F*_ST_ was developed using empirical Bayes (Beaumont and Balding 2004) and full Bayesian methods (BayeScan) (Foll and Gaggiotti 2008). The Bayesian methods have been applied in many cases of various species to identify outlier single-nucleotide polymorphisms (SNPs) (*e.g.*, Therkildsen *et al.* 2013a; Geraldes *et al.* 2014; Limborg *et al.* 2012). However, genome-wide population-specific *F*_ST_ is new among biologists. Despite the expected usefulness of genome-wide population-specific *F*_ST_ in evolutionary biology (Weir and Goudet 2017), applications have been sparse to date (*e.g.*, Nicholson *et al.* 2002; Weir *et al.* 2005; Foll and Gaggiotti 2006; Buckleton *et al.* 2016; Rougemont *et al.* 2020).

Traditional *F*_ST_ estimators were originally developed to estimate *F*_ST_ over a metapopulation (global *F*_ST_) based on a set of population samples (Cockerham 1969; Nei and Chesser 1983; Weir and Cockerham 1984; Excoffier 2007; Rousset 2007). In this study, we use Nei and Chesser’s (1983) bias-corrected *G*ST moment estimator (hereafter, NC83) as a pairwise *F*_ST_ estimator (Supplemental Note). The pairwise *F*_ST_ between populations (*i*, *j*) is defined as

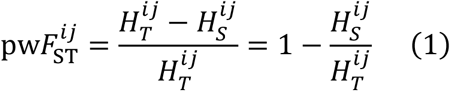

where 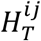 is total heterozygosity over all populations and 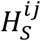 is within-population heterozygosity.

We apply Weir and Goudet’s (2017) bias-corrected moment estimator of population-specific *F*_ST_ (hereafter, WG) (Supplemental Note). When only allele frequencies are used, the WG population-specific *F*_ST_ at a locus is defined as

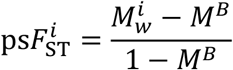

where 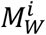 is the within-population matching of two distinct alleles of population *i* and *M*^B^ is the between-population-pair matching average over pairs of populations *i*, *i*′. *M*^B^ is homozygosity over pairs of populations. We represent heterozygosity over all pairs of populations as 1 - *M*^B^ = *H*_B_, and 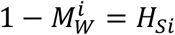. Therefore,

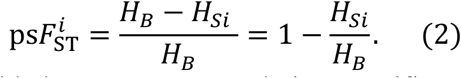

This formulation is reasonable because WG population-specific *F*_ST_ uses “allele matching, equivalent to homozygosity and complementary to heterozygosity as used by Nei (1973), rather than components of variance” (Weir and Goudet 2017). *H*_B_ is heterozygosity for all pairs of populations, whereas 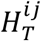 in Equation 1 is heterozygosity for the pair of populations. Equation 2 shows that WG population-specific *F*_ST_ measures population-specific genetic diversity (*H*_Si_) under the framework of the relatedness of individuals and identifies the population with the greatest genetic diversity as the ancestral or oldest population. Because populations close to the ancestral population have had more opportunities for mutations than recently founded populations (Liu *et al.*, 2006), they are likely to have the highest heterozygosity and low values of population-specific *F*_ST_. Thus, WG population-specific *F*_ST_ works to infer evolutionary history through genetic diversity in terms of heterozygosity under the assumption that the ancestral population had the highest genetic diversity. By combining population-specific and pairwise *F*_ST_ estimates, we can infer the present population structure, which reflects evolutionary history.

In this study, our objective is to demonstrate to empirical population geneticists and biologists how the two types of genome-wide *F*_ST_ estimators can be combined to help elucidate the population structure (pairwise *F*_ST_) in the evolutionary context (population-specific *F*_ST_). In our approach, the current population structure is represented by an unrooted neighbor-joining (NJ) tree (Saitou and Nei 1987) and a multi-dimensional scaling (MDS) plot based on pairwise *F*_ST_ values. The colors of the populations (names and/or sampling points) based on the WG genome-wide population-specific *F*_ST_ values enable the inference of the historical order of population colonization. We also present a representation on a geographical map, which is useful for visually understanding population history in a distribution range.

First, we examine the usefulness of our procedure using stepping-stone simulations that mimic population colonization from a single ancestral population for five scenarios of population range expansion. We then apply our approach to three datasets of human, Atlantic cod (*Gadus morhua*), and wild poplar (*Populus trichocarpa*). Human evolutionary history, migration, and population structure have been particularly well studied (*e.g*., Diamond 1997; Rosenberg *et al.* 2002; Ramachandran *et al.* 2005; Liu *et al.* 2006; Hellenthal *et al.* 2014; Rutherford 2016; Nielsen *et al.* 2017). These patterns are well known by statistical/theoretical population geneticists and biologists; therefore, testing our integrative *F*_ST_ analysis on this dataset could provide a good example of the usefulness of the practical approach. In humans, genome-wide SNPs have been used to infer population structure and admixture (Pickrell and Pritchard 2012; Lipson *et al.* 2013; Hellenthal *et al.* 2014; Bradburd *et al.* 2016). Conversely, microsatellite markers, which have a larger number of alleles than SNPs, despite the smaller number of loci, are still usual tools in ecology and conservation biology (*e.g*., Sylvester *et al.* 2018; D’Aloia *et al.* 2020; Kattenberg *et al.* 2020; Karamanlidis *et al.* 2021). We use microsatellite data for illustrative purposes. The Atlantic cod SNP were genotyped from mature fish samples collected from the North Atlantic from the northern range margin of the species in Greenland, Norway, and the Baltic Sea. Two ecotypes (migratory and stationary) that were able to interbreed were genetically differentiated (Hemmer-Hansen *et al.*, 2013a; Berg *et al.*, 2016). The inclusion of both types of data may help the understanding of the demographic history of highly migratory marine fish. The wild poplar SNP data were collected from the American Pacific Northwest. The wind-pollinated tree can spread pollen over a wide area, and the samples covered various regions over a range of 2,500 km near the Canadian–US border at altitudes between 0 and 800 m. Each sampling location was associated with 11 environmental and geographical parameters. The analysis of environmental variables and population-specific *F*_ST_ values may provide a good example for understanding the history of the range expansion of a wind-pollinated tree. By analyzing different types of data with species-specific ecology and migration history, the usefulness of our approach may be identified to enable us to understand the current population structure in an evolutionary context.

We computed pairwise and population-specific *F*_ST_ estimators using the R package FinePop2_ver.0.2, which is available at CRAN. The R codes for our representation method exemplified by the human data and simulations of population colonization used in this study are available in the Supplemental Material. They can be used for microsatellite and SNP genotype data in the Genepop format (Raymond and Rousset 1995; Rousset 2008), which has been particularly widely used among biologists.

## MATERIALS AND METHODS

### Population colonization simulations

To test the performance of our visual representation, we conducted simulations that mimicked the colonization of populations from a single ancestral population (population 1). We modeled five types of stepping-stone colonization: one, two, and three-directional population expansion; three-directional grid colonization from an edge; and eight-directional grid colonization from the center, with 24 demes (populations 2–25) (Figure 1A–E). We set the effective population size of the ancestral population to *N*_*e*_ = 10^4^ (twice the number of individuals in diploid organisms in a random mating population). At the beginning of colonization, 1% of *N*_*e*_ migrated into the adjacent vacant habitat once every 10 generations. For convenience, we considered one simulation cycle to be one generation. The effective population size of the newly derived population increased to the same size as the ancestral population (*N*_*e*_ = 10^4^) after one generation, and the population exchanged 1% of *N*_*e*_ genes with adjacent population(s) in every generation. Like the ancestral population, 1% of *N*_*e*_ individuals migrated into the adjacent vacant habitat once every 10 generations. We simulated the allele frequencies of SNPs in the ancestral and 24 derived populations. We also examined the cases in which the effective population size of the ancestral population was 10 times greater (*N*_*e*_ = 10^5^) than that of the newly derived population (*N*_*e*_ = 10^4^).

**Figure 1.**
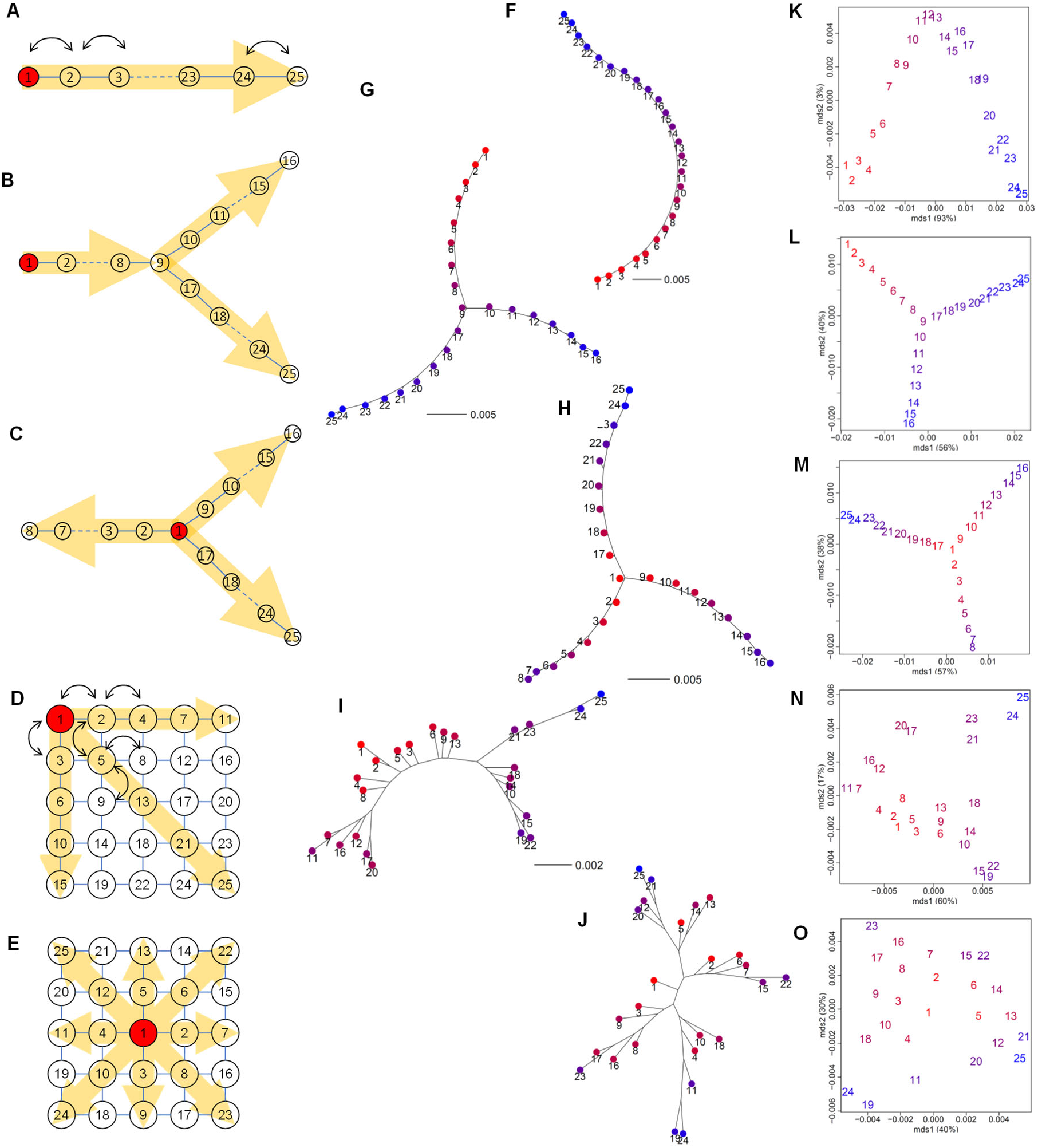
Results from population colonization simulations. Schematic diagrams of the models: (A) one, (B) two, (C) three-directional colonization, (D) three-directional grid colonization, and (E) eight-directional grid colonization. Population 1 in red is ancestral, and the yellow arrows indicate the direction of colonization. Lines show opportunities for migration. The effective population size of the newly derived population increased to the same size as the ancestral population (*N*_*e*_ = 10^4^) after one simulation generation, and each population exchanged 1% of *N*_*e*_ genes with adjacent population(s) in every generation, as indicated by the arrows (see the text). Neighbor-joining (NJ) unrooted trees (F–J) and multi-dimensional scaling (MDS) plots (K–O) based on the pairwise *F*_ST_ distance matrix overlaid with population-specific *F*_ST_ values for each model. The color of each population shows the magnitude of population-specific *F*_ST_ values between red (for the smallest *F*_ST_) and blue (for the largest *F*_ST_).

We generated the initial allele frequencies in the ancestral population, *q*, at 100,000 neutral SNP loci from the predictive equilibrium distribution, *f*(*q*) ∝ *q*^−1^(1 - *q*)^−1^ (Wright 1931). Additionally, we introduced 10 newly derived SNPs to each existing population in each generation. When a new SNP emerged in a population, we set the initial allele frequency of the newly derived SNP to 0.01 in the population and 0 in the other populations. This mimicked new mutations that survived in the initial phase after their birth. We considered these 100,000 ancestral SNPs and newly derived SNPs to be “unobserved.” We changed the allele frequencies of these SNPs using random drift under a binomial distribution in every generation. The frequencies of the derived alleles reduced for many of the SNPs over simulation generations and lost their polymorphism. After 260 simulation generations, we randomly selected SNPs that retained their polymorphism as “observed” SNPs. For grid colonization models (Figure 1D–E), we randomly selected polymorphic SNPs after 100 simulation generations. In this simulation, we selected 10,000 ancestral SNPs and 500 newly derived SNPs. Then, we generated 50 individuals for each population. We randomly generated the genotypes of these 10,500 SNPs for each individual following the allele frequencies in the population to which each individual belonged. Thus, we obtained “observed” genotypes of 1,250 individuals (= 50 individuals × 25 populations) at 10,500 SNP loci. To examine the effect of generations on genetic diversity in newly derived SNPs, we also selected 9,000 and 7,000 ancestral SNPs, and 1,000 and 3,000 newly derived SNPs, respectively. We converted the simulated genotypes into Genepop format (Raymond & Rousset, 1995; Rousset, 2008). We then computed genome-wide population-specific and pairwise *F*_ST_ values between the 25 populations.

### Visual representation of population structure and demographic history

We integrated genome-wide population-specific and pairwise *F*_ST_ estimates on a map of sampling locations on an NJ tree and MDS plot. We visualized the magnitude of the genome-wide population-specific *F*_ST_ values using a color gradient based on rgb (1 - *F*_ST,0_, 0, *F*_ST,0_), where *F*_ST,0_ = (*F*_ST_ - min*F*_ST_)/(max*F*_ST_ - min*F*_ST_). This conversion represents the standardized magnitude of a population-specific *F*_ST_ value at the sampling point, with colors between red (for the smallest *F*_ST_) and blue (for the largest *F*_ST_). We drew the *F*_ST_ map using the sf package in R, where we plotted sampling locations based on the longitudes and latitudes. We connected sampling points with genome-wide pairwise *F*_ST_ values smaller than a given threshold using yellow lines to visualize the connectivity between populations. Under the assumption of Wright’s island model at equilibrium between drift, mutation, and migration (Wright 1931), 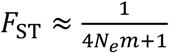, where *N*_*e*_ is the effective population size and *m* is the average rate of migration between populations (Slatkin 1987). For the threshold of *F*_ST_ = 0.02, there are 4*N*_*e*_*m* ≈ 49 migrants per generation (see Whitlock and Mccauley 1999; Waples and Gaggiotti 2006). We plotted the genome-wide population-specific *F*_ST_ values on a dot chart with standard errors estimated using Equation S5 (Supplemental Note). We drew the NJ tree based on the distance matrix of the genome-wide pairwise *F*_ST_ values using the `nj` function in the R package ape. We performed MDS analysis on the pairwise *F*_ST_ distance matrix using the `cmdscale` function in R. We used the cumulative contribution ratio up to the *k*th axis (*j* = 1, … , *k*, … , *K*) as the explained variation measure, which we computed using the R function as 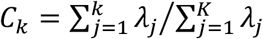, where *λ*_*j*_ is the eigenvalue and *λ*_*j*_ = 0 if *λ*_*j*_ < 0. We colored the sampling locations on the *F*_ST_ maps, dot charts, NJ trees, and MDS plots using a color gradient of the magnitude of genome-wide population-specific *F*_ST_ values. We also examined a diverging color palette instead of blue to red to test the resolution using RColorBrewer on the *F*_ST_ maps.

### Computing *F*_ST_ values

We converted the genotype data into Genepop format (Raymond and Rousset 1995; Rousset 2008) for implementation in the R package FinePop2_ver.0.2. We computed genome-wide pairwise *F*_ST_ values (NC83, Equation S3) using the `pop_pairwiseFST` function in FinePop2. We calculated expected heterozygosity for each population using the `read.GENEPOP` function. We computed the genome-wide WG population-specific *F*_ST_ (Equation S4) values using the `pop_specificFST` function. We applied Bayesian population-specific *F*_ST_ estimators on human data. We maximized Equation S7 and estimated the empirical Bayesian population-specific *F*_ST_ (Beaumont and Balding 2004) at each locus according to Equation S8. Then we averaged these values over all loci. For the full Bayesian model, we used GESTE_ver. 2.0 (Foll and Gaggiotti 2006) to compute the genome-wide population-specific *F*_ST_ values. We examined the shrinkage effect of the Bayesian population-specific *F*_ST_ estimator on inferring the ancestral population using a set of subsamples (37 populations) chosen from 51 populations.

### Inferring environmental effects on the observed population structure

To infer the geography and environment that were experienced by the population range expansion, we regressed the genome-wide population-specific *F*_ST_ values on the geographical and environmental variables (*j* = 1, … , *s*):

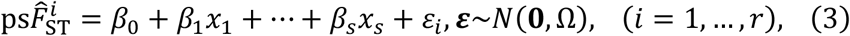

where Ω is the variance matrix of population-specific *F*_ST_. We correlated residuals because of the population structure; therefore, the effective sample size was lower than the actual sample size. In such circumstances, ordinary least squares overestimate the precision. To take the correlation into account, we derived the components of the variance–covariance matrix of the population-specific *F*_ST_ estimator for generalized least squares (GLS). We performed this analysis on the wild poplar dataset, for which 11 environmental/geographical parameters were available for each sampling location. We used Equations S5 and S6 for the components of the variance matrix Ω in Equation 3. We performed regression using the GLS function in FinePop2_ver.0.2.

### Three empirical datasets

We retrieved the human microsatellite data used in Rosenberg *et al.* (2002) from https://web.stanford.edu/group/rosenberglab/index.html. We removed the Surui sample (Brazil) from the data because that population was reduced to 34 individuals in 1961 as a result of introduced diseases (Liu *et al.* 2006). We retained genotype data (*n* = 1,035) of 377 microsatellite loci from 51 populations categorized into six groups, as in the original study: 6 populations from Africa, 12 from the Middle East and Europe, 9 from Central/South Asia, 18 from East Asia, 2 from Oceania, and 4 from America. We obtained the longitudes and latitudes of the sampling sites from Cann *et al.* (2002) (Table S1).

We combined the Atlantic cod SNP genotype data of 924 SNPs common to 29 populations reported in Therkildsen *et al.* (2013a, b) and 12 populations reported in Hemmer-Hansen *et al.* (2013a, b). We compared the genotypes associated with each marker in samples that were identical in the two studies, that is, CAN08 and Western_Atlantic_2008, ISO02 and Iceland_migratory_2002, and ISC02 and Iceland_stationary_2002, and standardized the gene codes. We removed cgpGmo.S1035, whose genotypes were inconsistent between the two studies. We also removed cgpGmo.S1408 and cgpGmo.S893, for which the genotypes were missing in several population samples in Therkildsen *et al.* (2013b). For simplicity, we removed temporal replicates from the Norway migratory, Norway stationary, North Sea, and Baltic Sea samples. The final dataset consisted of genotype data (*n* = 1,065) for 921 SNPs from 34 populations: 3 from Iceland, 25 from Greenland, 3 from Norway, and 1 each from Canada, the North Sea, and the Baltic Sea. All individuals in the samples were adults, and most were mature (Therkildsen *et al.*, 2013a). We used the longitudes and latitudes of the sampling sites in Hemmer-Hansen *et al.* (2013a). For the data from Therkildsen *et al.* (2013a), we estimated approximate sampling points from the map of the original study and recorded the longitudes and latitudes (Table S2).

We retrieved wild poplar SNP genotype data and environmental/geographical data from the original studies of McKown *et al.* (2014a, b). The genotype data contained 29,355 SNPs of 3,518 genes of wild poplar (*n* = 441) collected from 25 drainage areas (McKown *et al.*, 2014c). Details of the array development and selection of SNPs are provided in Geraldes *et al.* (2011, 2013). A breakdown of the 25 drainages (hereafter, populations) is as follows: 9 in northern British Columbia (NBC), 2 in inland British Columbia (IBC), 12 in southern British Columbia (SBC), and 2 in Oregon (ORE) (Geraldes *et al.*, 2014). We combined the original names of the clusters and population numbers, and used them as our population labels (NBC1, NBC3,…, ORE30). We associated each sampling location with 11 environmental and geographical parameters: latitude (lat), longitude (lon), altitude (alt), longest day length (DAY), frost-free days (FFD), mean annual temperature (MAT), mean warmest month temperature (MWMT), mean annual precipitation (MAP), mean summer precipitation (MSP), annual heat-moisture index (AHM), and summer heat-moisture index (SHM) (Table S3). The AHM was calculated in the original study as (MAT+10)/(MAP/1000); a large AHM indicates extremely dry conditions.

### Data availability

The authors affirm that all data necessary for confirming the conclusions of the article are present within the article, figures, a table, and supplemental material.

## RESULTS

### Simulations of population colonization

First, we examined the effect of the number of simulation generations on genetic diversity in newly derived SNPs using the eight-directional grid simulation (Figure 1E). WG population-specific *F*_ST_ correctly identified the ancestral population and traced the population history, and population structure reflected the population history regardless of the numbers of ancestral SNPs (9,000 and 7,000) and newly derived SNPs (1,000 and 3,000) selected after 100 simulation generations (Figures S1). The result was consistent with the case that used 10,500 SNP loci (10,000 ancestral SNPs + 500 newly derived SNPs) (Figures 1J, O, S2E). In the following analysis, we used the results based on 10,500 SNP loci to generate clearer results, even for limited numbers of simulation generations (260 and/or 100).

In the one-directional simulation (Figure 1A), population-specific *F*_ST_ correctly identified the ancestral population with the highest genetic diversity (Figure S2A), and populations were located in order from 1 to 25 on the NJ tree (Figure 1F). The first axis of the MDS plot explained 93% of the variance of the pairwise *F*_ST_ distance matrix and indicated population expansion from population 1 to population 25 (Figure 1K). In the two-directional simulation (Figure 1B), our analysis correctly identified the ancestral population (Figure S2B) and detected that populations were split at population 9 and expanded in two directions (Figure 1G), which was consistent with the simulation scenario. The first axis of the MDS plot identified population expansion from population 1 to population 25 and explained 56% of the variance of the pairwise *F*_ST_ distance matrix, whereas the second axis identified another manner of population expansion from population 1 to population 16 and explained 40% of the variation (Figure 1L). In the three-directional simulation (Figure 1C), the ancestral population was also correctly identified (Figure S2C). It was closely located to the adjacent populations 2, 9, and 17, but correctly detected three directions (Figure 1H). The first axis of the MDS plot identified population expansion from population 1 to populations 16 and 25, and explained 57% of the variance of pairwise *F*_ST_, whereas the second axis identified population expansion from population 1 to populations 8 and 16 and explained the 38% variation (Figure 1M).

In the three-directional grid colonization model from an edge (Figure 1D), population-specific *F*_ST_ correctly identified the ancestral population (Figure S2D) and pairwise *F*_ST_ detected that populations expanded in three directions (Figure 1I), which agreed with the simulation scenario. The first axis of the MDS plot identified population expansion from population 1 to other edge populations (populations 11, 15, and 25), and explained 60% of the variance of the pairwise *F*_ST_ distance matrix, whereas the second axis indicated genetic differentiation between populations 24, 25 and 15, 19, 22, and explained 17% of the variation (Figure 1N). In the eight-directional grid colonization model (Figure 1E), population-specific *F*_ST_ identified the ancestral population (Figure S2E) and pairwise *F*_ST_ estimated that populations expanded in five directions from the center (Figure 1J). The first axis of the MDS plot identified vertical population expansion from population 1 to populations 24 and 25 and explained 40% of the variance of the pairwise *F*_ST_ distance matrix, and the second axis indicated horizontal population expansion from population 1 to populations 24 and 23, which explains 30% of the variation (Figure 1O). We obtained similar results in the cases in which the effective population size of the ancestral population was 10 times greater (*N*_*e*_ = 10^5^) than that of the newly derived population (*N*_*e*_ = 10^4^) (Figure S3). We obtained very similar results from different data for more than 20 simulations (figures not shown).

### Humans

The *F*_ST_ map (Figure 2A) shows integrated information from genome-wide population-specific and pairwise *F*_ST_, which visualizes population structure with the migration and demographic history of human populations in terms of genetic diversity. Interestingly, Bantu Kenyans had the smallest *F*_ST_ value (shown in red). Figure 2B ordered population-specific *F*_ST_ values from Africa to Central/South Asia, the Middle East, Europe, East Asia, Oceania, and America (Table S4). As indicated by the sampling points connected by yellow lines with pairwise *F*_ST_ values below the 0.02 threshold (Figure 2A), genetic connectivity from Africa was low. Conversely, migration was substantial within Eurasia but much smaller than that inferred from Eurasia to Oceania and America. *H*_*e*_ was the highest in Africa, followed by the Middle East, Central/South Asia, Europe, and East Asia, but relatively small in Oceania and lowest in South America. The Karitiana in Brazil had the lowest *H*_*e*_. The NJ tree (Figure 2C) integrated with population-specific *F*_ST_ values indicated that human populations originated from Bantu Kenyans and expanded to Europe through Mozabite, the Middle East, Central/South Asia, and East Asia. The Kalash was isolated from Europe/Middle East and Central/South Asia populations. Papuans/Melanesians and American populations diverged from between Central/South Asian and East Asian populations. The ordinal NJ tree of pairwise *F*_ST_ values divided the populations into five clusters: 1) Africa, 2) the Middle East, Europe, and Central/South Asia, 3) East Asia, 4) Oceania, and 5) America (Figure S4). The first axis of the MDS plot highlighted difference between African and American populations and explained 44% of the variance of the pairwise *F*_ST_ distance matrix, whereas the second axis indicated genetic differentiation between Melanesian and Karitiana populations, and explained the 19% variation (Figure 2D).

**Figure 2.**
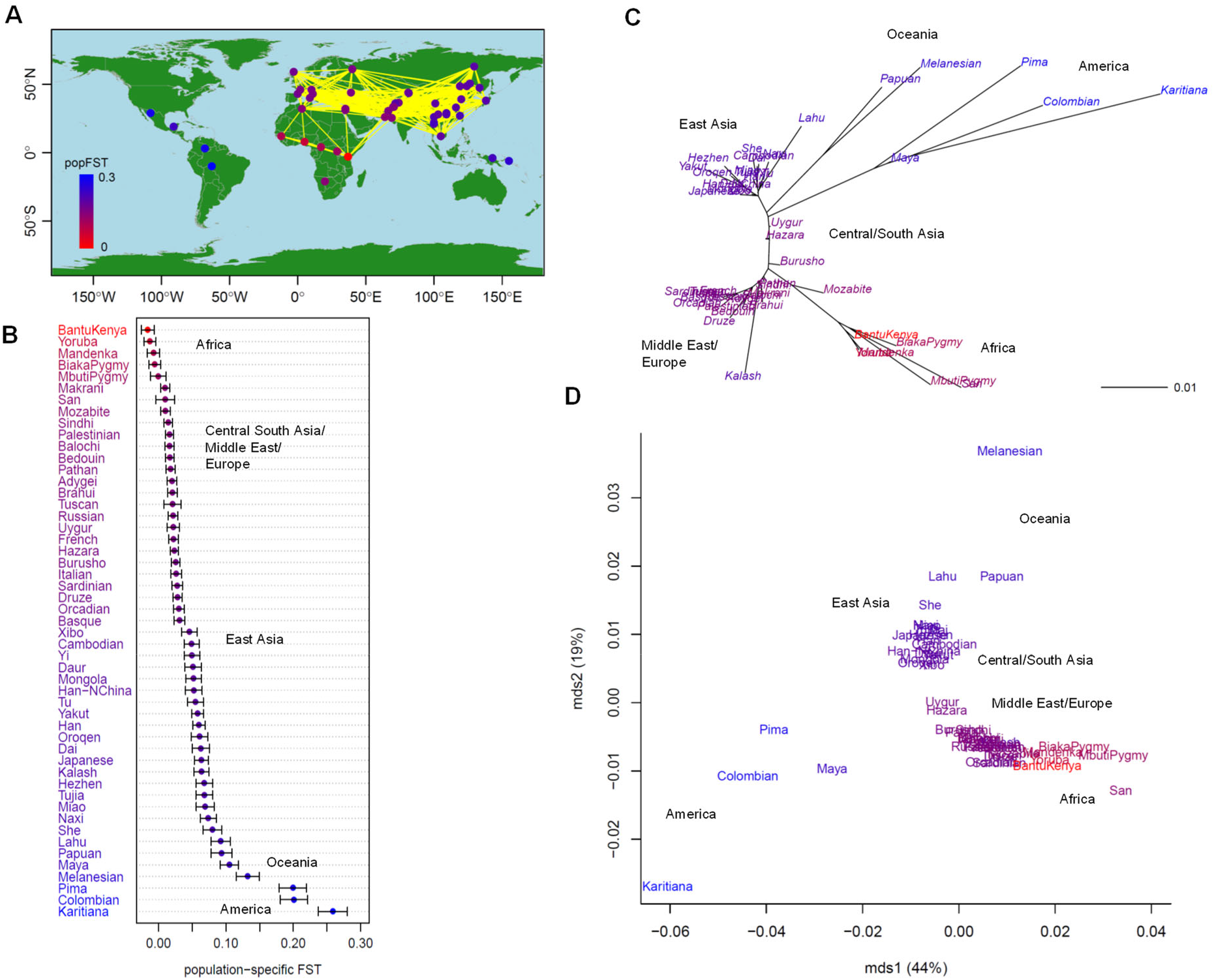
Population structure of 51 human populations (*n* = 1,035; 377 microsatellites). (A) Map showing population connectivity with the magnitude of population-specific *F*_ST_ values. Populations connected by yellow lines are those with pairwise *F*_ST_ < 0.02. (B) Distribution of population-specific *F*_ST_ values ± 2×SE. (C) Neighbor-joining (NJ) unrooted tree and (D) multi-dimensional scaling (MDS) based on pairwise *F*_ST_ overlaid with population-specific *F*_ST_ values on population labels. The color of each population indicates the magnitude of population-specific *F*_ST_ values between red (for the smallest *F*_ST_) and blue (for the largest *F*_ST_).

The Bayesian population-specific *F*_ST_ values estimated using the methods of Beaumont and Balding (2004) (empirical Bayes) and Foll and Gaggiotti (2006) (full Bayes) were nearly identical and the smallest *F*_ST_ values observed in the Middle East, Europe, and Central/South Asia (Figure S5A, Table S4). However, in African populations, they were higher than the WG population-specific *F*_ST_ values (Figure S5B). Our *F*_ST_ map based on the empirical Bayesian population-specific *F*_ST_ values indicated that the Middle East, Europe, and Central/South Asia were centers of human origin (Figure 3A), which was consistent with that from the full Bayesian population-specific *F*_ST_ estimator (figure not shown). Our integrated NJ tree showed that the Hazara, Pakistan population was genetically closest to the human ancestors (Figure S6A). The numbers of sampling locations of the 51 human populations were as follows: 6 from Africa, 12 from the Middle East/Europe, 9 from Central/South Asia, 18 from East Asia, 2 from Oceania, and 4 from America. When we used a subsample of the 37 human populations (3 from Africa, 6 from the Middle East/Europe, 4 from Central/South Asia, 18 from East Asia, 2 from Oceania, and 4 from America; Table S5), the area with the highest population-specific *F*_ST_ values shifted toward Central/South Asia and East Asia (Figure 3B), whereas Bantu Kenyans had the smallest WG population-specific *F*_ST_ value (Figure 3C); this was consistent with the results from the full dataset (Figure 2A). The integrated NJ trees provided similar results (Figure S6B, C).

**Figure 3.**
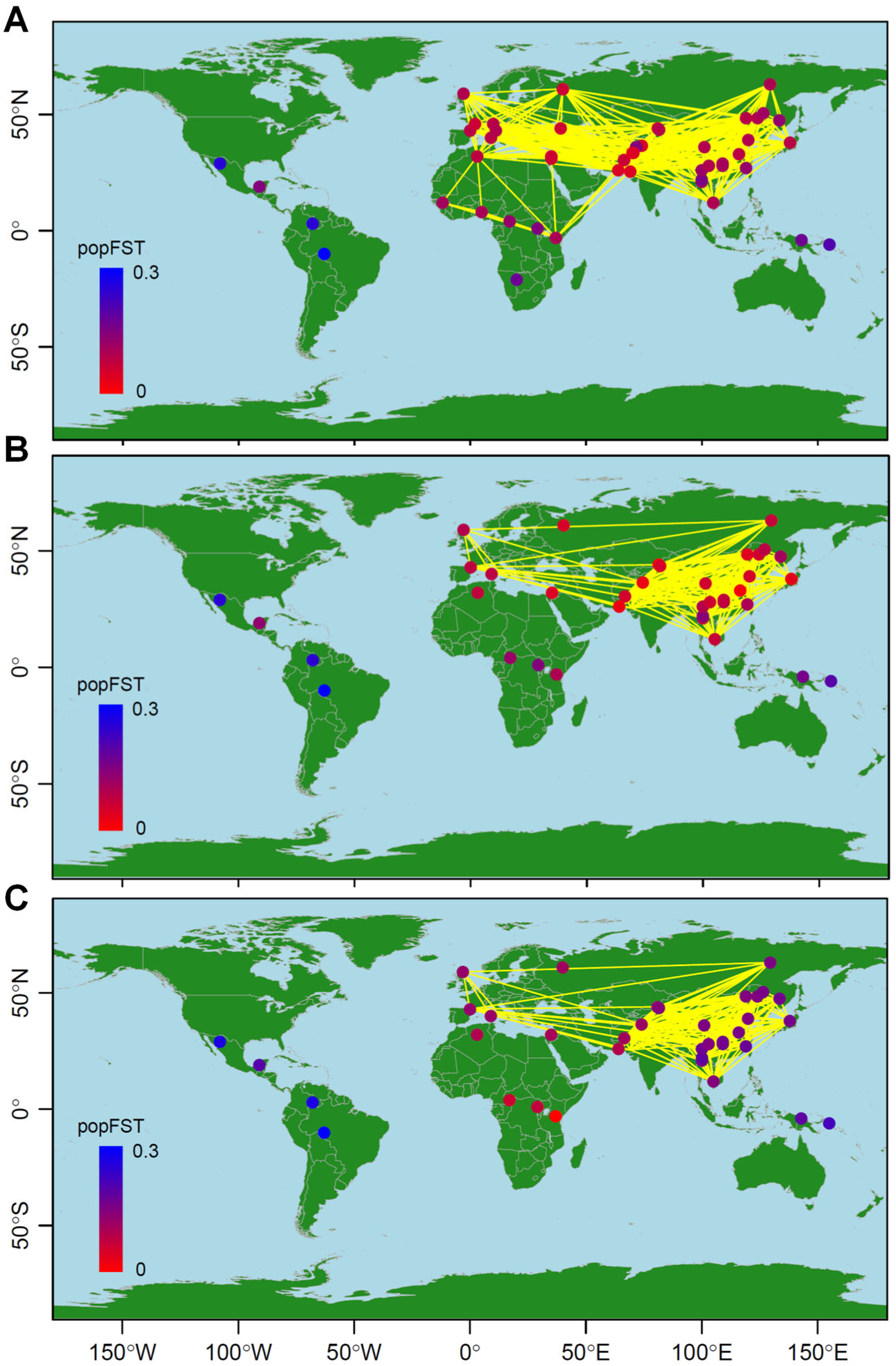
Population structure of humans based on Bayesian and moment estimators of population-specific *F*_ST_. Results from the Bayesian population-specific *F*_ST_ estimator using (A) 51 samples and (B) 37 subsamples, and from (C) the WG population-specific *F*_ST_ moment estimator using 37 subsamples. The numbers of sampling locations of the subsamples were as follows: 3 from Africa, 6 from the Middle East/Europe, 9 from Central/South Asia, 18 from East Asia, 2 from Oceania, and 4 from America. Populations connected by yellow lines are those with pairwise *F*_ST_ < 0.02. The color of each population indicates the magnitude of population-specific *F*_ST_ values between red (for the smallest *F*_ST_) and blue (for the largest *F*_ST_).

### Atlantic cod

The *F*_ST_ map (Figure 4A) visualizes the population structure, migration, and genetic diversity of the Atlantic cod populations. The Canadian population had the smallest population-specific *F*_ST_ value (shown in red) and the greatest *H*_*e*_. *H*_*e*_ was also high in Greenland, low in other areas, and lowest in the Baltic Sea. Figure 4B shows the order of population-specific *F*_ST_ values from Canada to the Baltic sea (Table S6). Greenland west coast populations (green in Figure S7) generally had small population-specific *F*_ST_ values, whereas fjord populations (violet) had relatively higher population-specific *F*_ST_ values. The population-specific *F*_ST_ values were much higher for populations in Iceland, Norway, and the North Sea, and highest in the Baltic Sea. Based on pairwise *F*_ST_ values between sampling points (< 0.02 threshold) (Figure 4A), substantial migration was suggested between Greenland, Iceland, and Norway. Conversely, migration could be low from Canada to Greenland and from Iceland and Norway to the North and Baltic Seas. Our integrated NJ tree with population-specific *F*_ST_ values (Figure 4C) inferred that Atlantic cod originated from Canada, migrated to the west coast of Greenland, and then expanded their distribution to Iceland, Norway, the North Sea, and the Baltic Sea. According to the ordinal NJ tree of the pairwise *F*_ST_ distance matrix (Figure S7), the populations were divided into four large clusters: 1) Canada; 2) Greenland west coast, 3) Greenland east coast, Iceland, and Norway; and 4) North and Baltic Seas. Fjord populations (in purple) formed a sub-cluster within the Greenland west coast, and migratory (orange) and stationary (magenta) ecotypes also formed a sub-cluster. The first axis of the MDS plot characterized the differentiation between Canadian and North Sea/Baltic Sea populations and explained 72% of the variance of the pairwise *F*_ST_ distance matrix, whereas the second axis highlighted the differentiation between Norwegian migratory type populations from Canadian and North Sea/Baltic Sea populations, which explains the 22% variation (Figure 4D).

**Figure 4.**
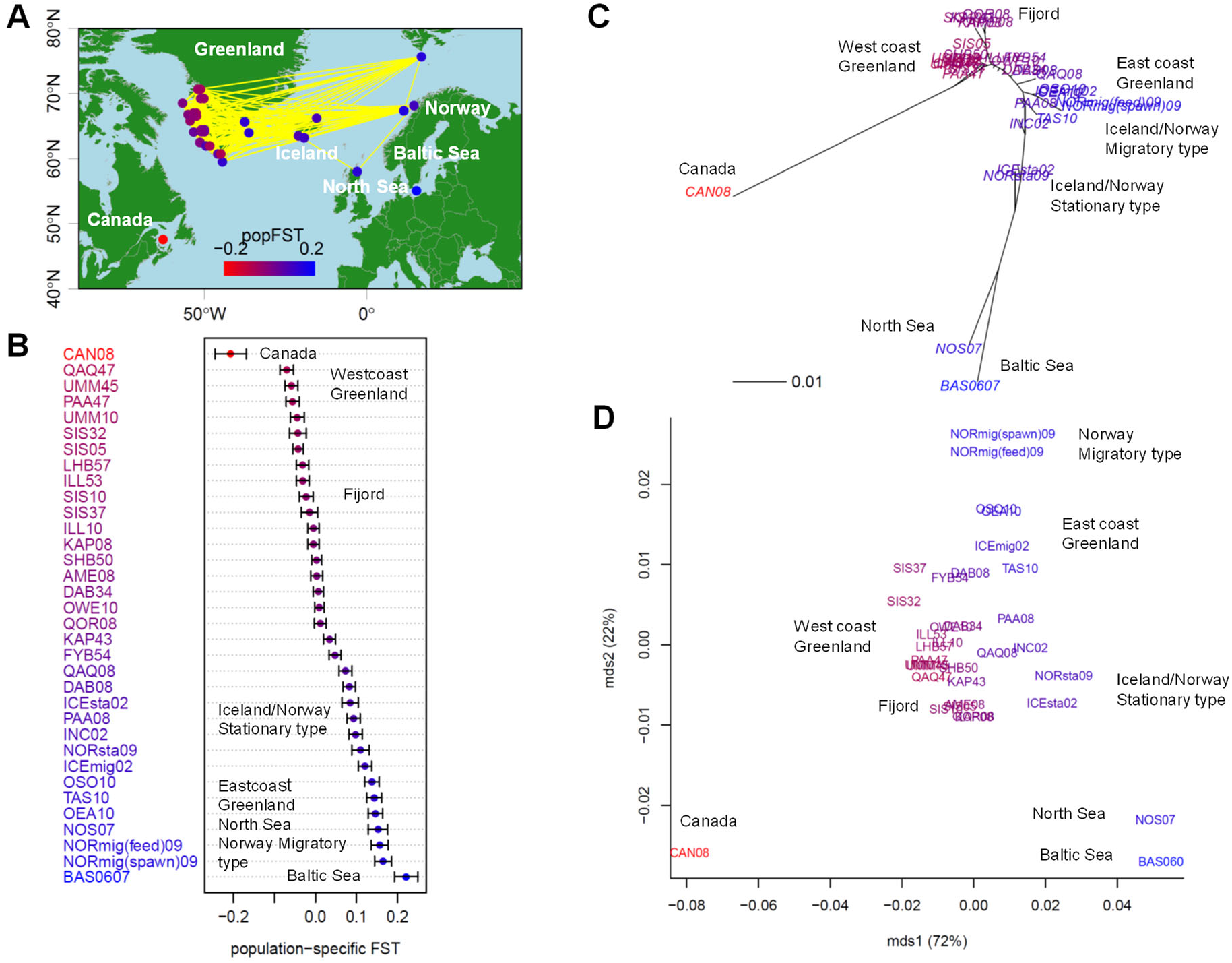
Population structure of 34 geographical samples of wild Atlantic cod (*n* = 1,065; 921 SNPs). (A) Map showing population connectivity with the magnitude of population-specific *F*_ST_ values. Populations connected by yellow lines are those with pairwise *F*_ST_ < 0.02. (B) Distribution of population-specific *F*_ST_ values ± 2×SE. (C) Neighbor-joining (NJ) unrooted tree and (D) multi-dimensional scaling (MDS) based on pairwise *F*_ST_ overlaid with population-specific *F*_ST_ values on population labels. The color of each population shows the magnitude of population-specific *F*_ST_ values between red (for the smallest *F*_ST_) and blue (for the largest *F*_ST_).

### Wild poplar

The *F*_ST_ map (Figure 5A) indicated that population-specific *F*_ST_ values were lowest in southern British Columbia (SBC27) and inner British Columbia (IBC15 and IBC16) (shown in red, Figure S8). The sampling points connected by yellow lines (< 0.02 threshold pairwise *F*_ST_ values) indicated migration between all populations. *H*_*e*_ was highest in SBC27, IBC15, and IBC16, and lowest in northern British Columbia (NBC5). Figure 5B shows that samples collected from areas close to the SBC coast had higher population-specific *F*_ST_ values than other SBC samples (Table S7). The NBC samples had population-specific *F*_ST_ values similar to those of SBC. Among the NBC samples, NBC8 had the smallest population-specific *F*_ST_, and NBC5 had the highest value, followed by NBC6 and NBC7. The pairwise *F*_ST_ NJ tree integrated with population-specific *F*_ST_ values (Figure 5C) suggested that wild poplar originated from around SBC27 and Inner British Columbia, and expanded in three directions: to the BC southern coast, northern BC and south-western Alaska, and Oregon. The ordinal NJ tree based on the pairwise *F*_ST_ distant matrix divided populations into four large clusters: 1) IBC, 2) SBC, 3) NBC, and 4) ORE (Figure S8). The population represented by sample ORE30 was isolated from ORE29. The first axis of the MDS plot characterized the differentiation between southern and northern populations and explained the 51% variance of the pairwise *F*_ST_ distance matrix, whereas the second axis characterized the southernmost ORE30 population, and explained the 18% variation (Figure 5D).

**Figure 5.**
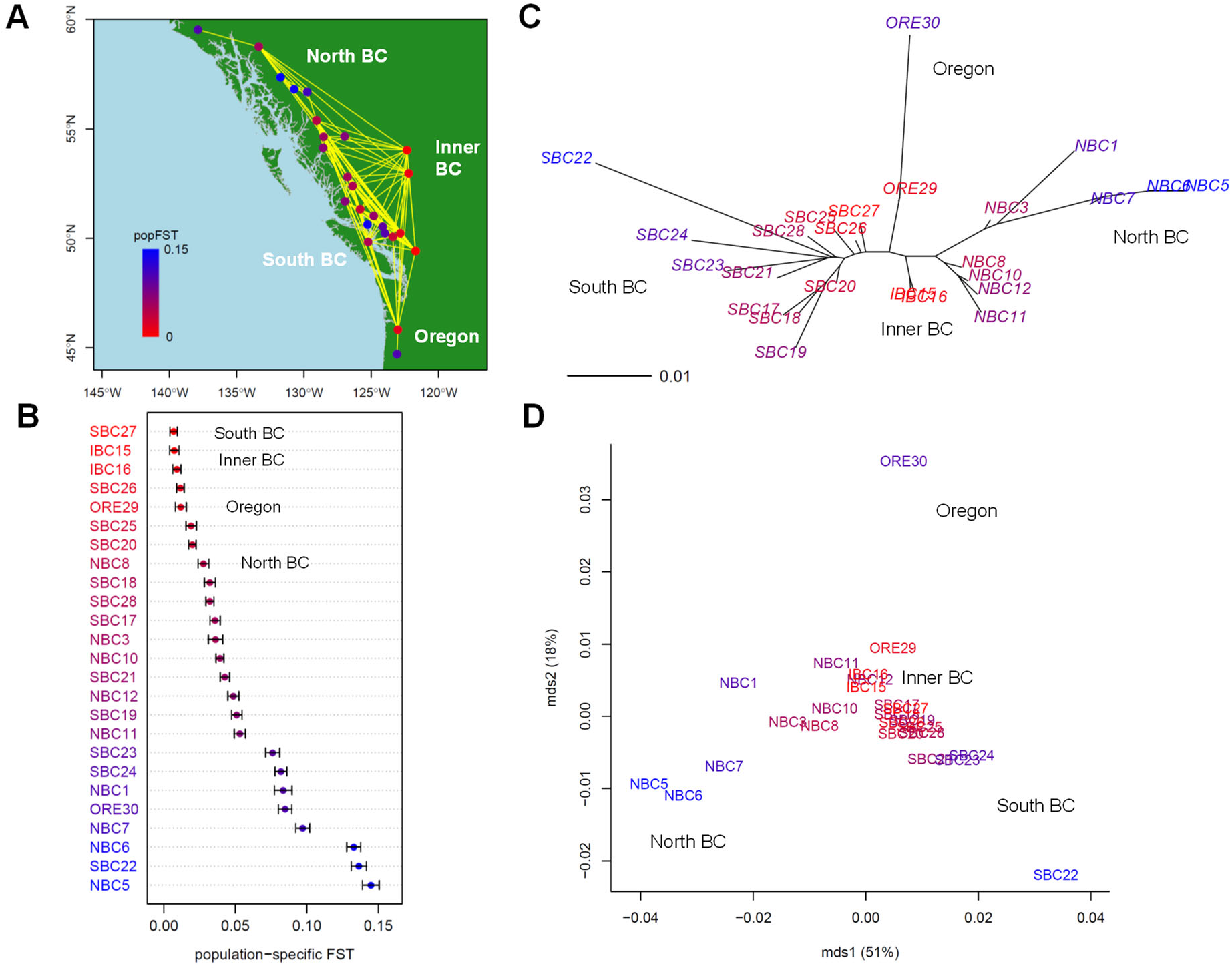
Population structure for 25 geographical samples of wild poplar (*n* = 441; 29,355 SNPs). (A) Map showing population connectivity with the magnitude of population-specific *F*_ST_ values. Populations connected by yellow lines are those with pairwise *F*_ST_ < 0.02. (B) Distribution of population-specific *F*_ST_ values ± 2×SE. (C) Neighbor-joining (NJ) unrooted tree and (D) multi-dimensional scaling (MDS) based on pairwise *F*_ST_ overlaid with population-specific *F*_ST_ values on population labels. The color of each population indicates the magnitude of population-specific *F*_ST_ values between red (for the smallest *F*_ST_) and blue (for the largest *F*_ST_).

To avoid multicollinearity, we excluded seven out of 11 environmental variables that were significantly correlated with each other: lat, lon, alt, FFD, MWMT, MSP, and AHM (Table S3). Our GLS of genome-wide population-specific *F*_ST_ values on the four environmental variables (DAY, MAT, MAP, and SHM) indicated that DAY, MAP, and SHM were significant (Table 1). All estimates were positive, which indicated that higher population-specific *F*_ST_ values were expected for longer DAY (longer daylight time), higher MAP (abundant rain), and higher SHM (dry summers), and these values might reflect the directions of population expansion. The scatter plot of DAY and SHM (each population colored by the population-specific *F*_ST_ value) (Figure 6A) suggested three directions of population range expansion; the wild poplar that might originate from around SBC27 and IBC15 expanded its distribution to NBC, where daylight hours are long in summer; expanded to SBC seashores with lower SHM (humid summer and abundant rainfall), and to the south (ORE29 and ORE30) with higher SHM (dry summer). This was consistent in the scatter plot of DAY and MAP (Figure 6B).

**Table 1.** Regression of genome-wide population-specific *F*_ST_ of 25 wild poplar populations on environmental variables

| Variable | Estimate | SE | Z | *p* |
| --- | --- | --- | --- | --- |
| DAY | 0.0489 | 0.0164 | 2.99 | 0.003** |
| MAT | -0.0088 | 0.0086 | -1.03 | 0.305 |
| MAP | 0.0001 | 0.0000 | 2.79 | 0.005** |
| SHM | 0.0022 | 0.0009 | 2.38 | 0.018* |

DAY; longest day length (hours), MAT; mean annual temperature (°C), MAP; mean annual precipitation (mm), SHM; summer heat-moisture index, \* $p < 0.05$ and \*\* $p < 0.01$

**Figure 6.**
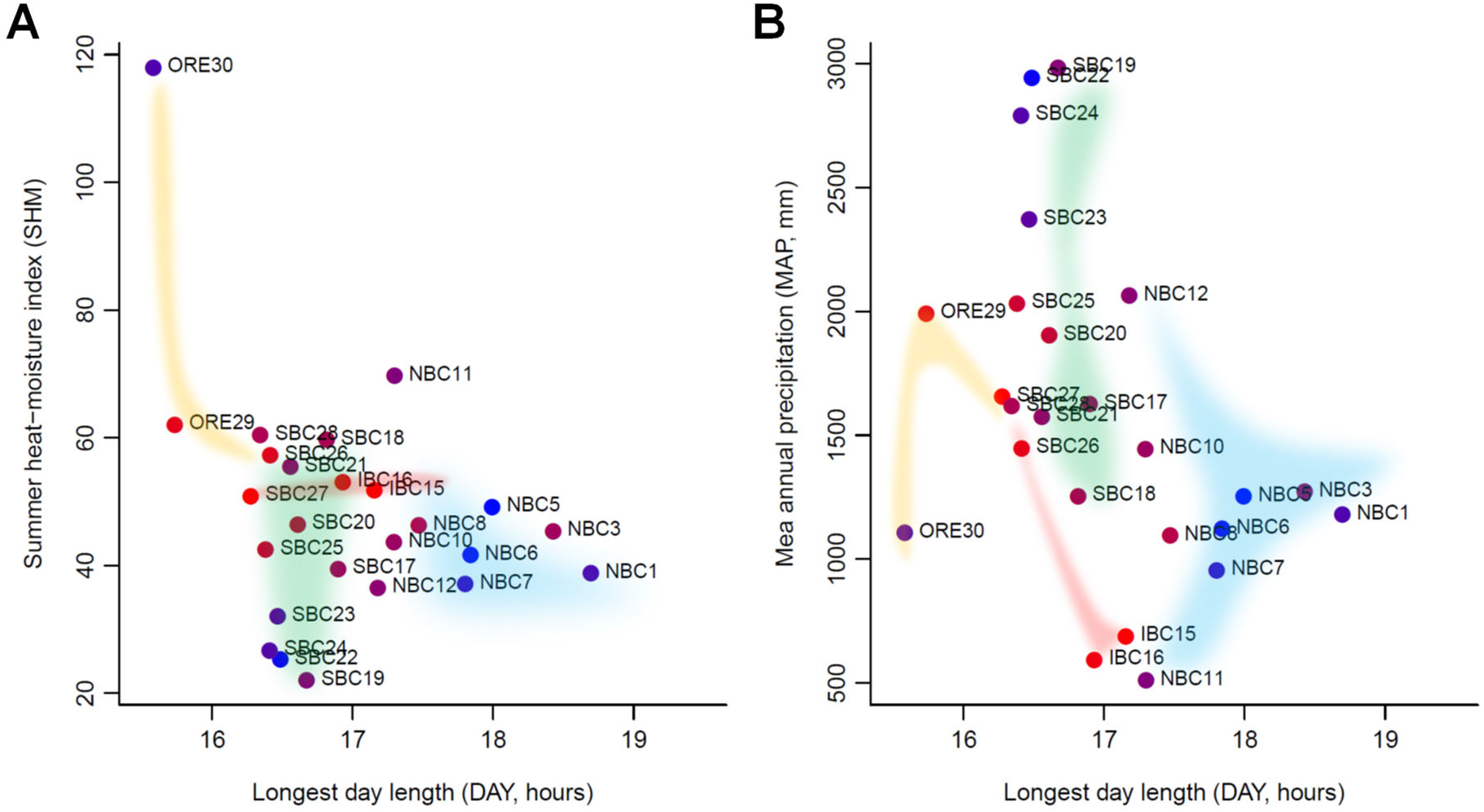
Population range expansion and key environmental variables. Longest day length vs. (A) summer heat-moisture index and (B) mean annual precipitation for 25 geographical samples of wild poplar. The colored areas by the population clusters (see Figure S8) show the inferred population expansion from IBC15, IBC16, and SBC27. The color of each population shows the magnitude of population-specific *F*_ST_ values between red (for the smallest *F*_ST_) and blue (for the largest *F*_ST_).

### Diverging color palette

RColorBrewer had 35 color palettes. Each palette had a minimum of eight colors. Two palettes had 12 colors (maximum) and nine had 11 colors. We chose a color palette with 10 colors (RdYlBu). The color gradient better identified the middle range of population-specific *F*_ST_ values, but failed to detect the ancestral population (Figure S9).

### CPU times

Using a laptop computer with an Intel Core i7-8650U CPU, only 89.8 s of CPU time was required to compute the WG population-specific *F*_ST_ estimates and SEs of wild poplar (29,355 SNPs; 25 populations, *n* = 441). Alternatively, 120.7 s was required to obtain the pairwise *F*_ST_ (NC83) between all population pairs. Based on these results, 50 min may be required to compute WG population-specific *F*_ST_ and 70 min to compute pairwise NC83 *F*_ST_ estimates for 1 million SNPs using that laptop. This computation may be much faster if we execute it on a workstation.

## DISCUSSION

### Genome-wide population-specific *F*_ST_ traced population history as reflected by genetic diversity

Our simulations demonstrated that the WG population-specific *F*_ST_ estimator identified the source population and traced the evolutionary history of its derived populations based on genetic diversity (heterozygosity estimated from each population). The NC83 pairwise *F*_ST_ estimator correctly estimated the current population structure. As explained in the Introduction, the population-specific *F*_ST_ estimator is a rescaling of expected heterozygosity, and we expect a linear relationship between expected heterozygosity and population *F*_ST_. This shows that the population-specific *F*_ST_ estimator implicitly assumes that populations closest to the ancestral population have the highest heterozygosity. In our three case studies, a linear relationship between *H*_*e*_ of each population (= *H*_Si_) and 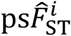 was evident (Figure S10). The coefficient of determination, *R*^2^, was 0.91 for 51 human populations (*n* = 1,035), 0.99 for 34 Atlantic cod populations (*n* = 1,065), and 0.82 for 25 wild poplar populations (*n* = 441). The goodness of fit to the linear function should depend on the sample size (number of individuals). Our simulations evaluated the performance of the population-specific *F*_ST_ estimator for such cases. However, in populations that experienced extensive admixture events, heterozygosity was enhanced, whereas a bottleneck in the ancestral population reduced heterozygosity. In such cases, the population-specific *F*_ST_ estimator misidentifies the ancestral population.

In our analysis, the genome-wide WG population-specific *F*_ST_ values successfully illustrated human evolutionary history, and indicated that humans originated in Kenya, expanded from the Middle East into Europe and from Central/South Asia into East Asia, and then possibly migrated to Oceania and America (Figure 2). Kenya is located just below Ethiopia, where the earliest anatomically modern humans were found from fossils (Nielsen *et al.* 2017). Our results are also in good agreement with the highest levels of genetic diversity being detected in Africa (Rosenberg *et al.*, 2002), the relationship uncovered between genetic and geographic distance (Ramachandran *et al.*, 2005), the shortest colonization route from East Africa (Liu *et al.*, 2006), and major migrations inferred from genomic data (Nielsen *et al.*, 2017).

Our analysis indicated that Atlantic cod might originate in Canada (CAN08). Figure 4 suggested that the population expansion of Atlantic cod began by minimal gene flow from Canada. They might have first expanded to the west coast of Greenland before spreading to Iceland, the North Sea, Norway, and the Baltic Sea. This result was consistent with genomic evidence that Atlantic cod inhabit both sides of the Atlantic Ocean and evolved from a common evolutionary origin (Berg *et al.* 2017). The migratory ecotypes characterized by deeper and more offshore habitats and long-distance migrations (Hemmer-Hansen *et al.*, 2013a) may have played an important role in this expansion, as suggested in Figures 4C and S7. In our study, CAN08 had the highest *H*_*e*_, which was lower in Iceland than in Greenland; this result implies that Icelandic populations were the descendants of colonists from Greenland, which in turn originated in Canada. The BAS0607 sample from the Baltic Sea had the highest population-specific *F*_ST_ and the lowest *H*_*e*_, suggesting that Baltic cod is the newest population. This result agrees with the findings of a previous study, which identified Baltic cod as an example of a species subject to ongoing selection for reproductive success in a low salinity environment (Berg *et al.* 2015). In the Atlantic cod case study, CAN08 had the highest *H*_*e*_ and a very large negative population-specific *F*_ST_ value of −0.21 ± 0.019 compared with the maximum value of 0.22 ± 0.014 in BAS0607 (Figure S10, Table S6). The WG population-specific *F*_ST_ value can be negative (Weir and Goudet 2017). In the one and two-directional models of our simulations, the WG population-specific *F*_ST_ value was significantly negative in the ancestral population, whereas *H*_*e*_ was the largest (Figure S2A,B). Our consistent results between the simulations and Atlantic cod case study indicate that when gene flow from other populations into the source population is limited, a relatively large 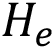 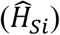 is maintained in the source population. In such cases with 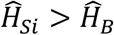, Equation 2 produces negative values for population-specific *F*_ST_.

Although the wild poplar samples used in this study might not cover the entire distribution range of the species, which extends from southern California to northern Alaska, Montana, and Idaho (Geraldes *et al.* 2013), the genome-wide population-specific *F*_ST_ values suggested three directions of population expansion of wild poplar: from southern British Columbia (SBC27) and inland British Columbia (IBC15, IBC16) to coastal British Colombia, southern Oregon, and northern British Columbia (Figure 5). The largest population-specific *F*_ST_ value was found in the population with the smallest heterozygosity, SBC22, which may have resulted from a bottleneck (Geraldes *et al.*, 2014).

Our continuous color gradient from red to blue successfully detected the ancestral population, whereas the R diverging color palettes, which had a limited number of colors (maximum of 12), better identified the middle range of population-specific *F*_ST_ values (Figure S9). Both color gradients may be useful.

The two types of genomewide *F*_ST_s enabled us to infer the current population structure and the history of range expansion, which can also be calculated for the windows in the genomes. By comparing the pairwise *F*_ST_ among windows, it is possible to identify the genomic regions that are largely different between the populations (*e.g.*, Yi *et al.* 2010). Likewise, by comparing population-specific *F*_ST_ among windows, it may be possible for us to identify unique populations and their corresponding unique genomic regions.

### Genome-wide population-specific *F*_ST_ detects key environments that relate to population expansion

Our GLS of genome-wide population-specific *F*_ST_ values revealed that long daylight hours, abundant rainfall, and dry summer conditions are the key environmental factors that relate to the demographic history of wild poplar (Table 1). Wild poplar could have expanded its distribution by its fluffy seeds being blown away by the wind. In the NJ unrooted tree, the root of the populations cannot be fixed without out-group populations. There is no direct evidence, but we can infer from the population-specific *F*_ST_ values that wild poplar seems to have spread from southern British Columbia (SBC27) and inland British Columbia (IBC15, 16), and expanded its distribution to NBC, where daylight hours are long in summer, to SBC seashores with the rainy environment, and to southern Oregon (ORE30) with dry summer conditions (Figures 5,6). A previous study on wild poplar revealed that genes involved in drought response were identified as *F*_ST_ outliers using Bayescan (Foll and Gaggiotti 2008) (Geraldes *et al.* 2014). The *F*_ST_ outlier test of Geraldes and colleagues also revealed that genes involved in the circadian rhythm and response to red/far-red light had high locus-specific global *F*_ST_ values. The first principal component of SNP allele frequencies of the polar tree was significantly correlated with day length, and a previous enrichment analysis for population structuring uncovered genes related to the circadian rhythm and photoperiod (McKown *et al.* 2014a). Our results were in agreement with previous findings, showing the usefulness of using the GLS of genome-wide population-specific *F*_ST_ to infer environmental effects on the population expansion of species.

### Genome-wide *F*_ST_ moment estimators converge to their true means

Previous studies have suggested or indicated that the “ratio of averages” works better than the “average of ratios” as the number of independent SNPs increases (Reynolds *et al.* 1983; Weir and Cockerham 1984; Bhatia *et al.* 2013). Because “the combined ratio estimate (ratio of averages) is much less subject to the risk of bias than the separate estimate (average of ratios)” (Cochran 1977), scholars recommend using the “ratio of averages” estimators (Bhatia *et al.* 2013). To explicitly show the underlying mechanism, we used the observed heterozygosity of population 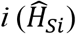 as derived by Nei and Chesser (1983) (Supplemental Note). When the number of loci (*L*) increases, the average observed heterozygosity over all loci converges to its expected value according to the law of large numbers as

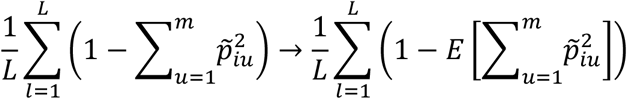

The observed gene diversity thus converges to the expected value:

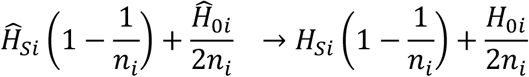

Similarly, 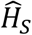 and 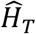 converge to their expected values. This example indicates that the numerators and denominators of bias-corrected *F*_ST_ moment estimators, whether global, pairwise, or population-specific, converge to their true means and provide unbiased estimates of *F*_ST_ in population genomics analyses with large numbers of SNPs.

### Bayesian *F*_ST_ estimators measure the deviation from the average of the sampled populations

In the Bayesian framework, the population specific *F*_ST_ is the coefficient of the genetic drift that represent the among-population variation of the allele frequencies at neutral loci from the allele frequencies of the ancestral population. The allele frequency of the ancestral population is assumed to be the among-population mean allele frequencies. The analysis of human population (Figure 3A) provokes the need to take account of geographical heterogeneity of sampling fraction of populations. The unbiased estimate of the allele frequency of the ancestral population will be obtained as the weighted among-population average. The weights are inversely proportional to the sampling fractions.

The shrinkage effect on allele frequencies in Bayesian inference (Stein, 1956) may shift population-specific *F*_ST_ values toward the average of the entire population. Because of the shrinkage toward mean allele frequencies, the maximum likelihood and Bayesian estimators of locus-specific global *F*_ST_ improve the power to detect genes under environmental selection (Beaumont and Balding 2004; Foll and Gaggiotti 2008). An empirical Bayes genome-wide pairwise *F*_ST_ estimator (Kitada *et al.* 2007) is useful in cases involving a small number of polymorphic marker loci, particularly in high gene flow scenarios, but it suffers from the shrinkage effect when larger numbers of loci are used. The shrinkage of allele frequencies should affect inference in genome-wide population-specific *F*_ST_, particularly in cases when samples (populations) were not a representative of the populations.

### Conclusions

The WG genome-wide population-specific *F*_ST_ moment estimator can identify the source population and trace the evolutionary history of the derived populations based on genetic diversity under the assumption that populations closest to the ancestral population have the highest heterozygosity. Conversely, the NC83 genome-wide pairwise *F*_ST_ moment estimator represents the current population structure. By integrating population-specific and pairwise estimates on *F*_ST_ maps, NJ trees, and MDS plots, we obtain a picture of population structure by incorporating evolutionary history. Our GLS analysis of genome-wide population-specific *F*_ST_, which takes the correlation between populations into account, provides insights into how a species expanded its distribution in different environments. Given a large number of loci, bias-corrected *F*_ST_ moment estimators – whether global, pairwise, or population-specific – provide unbiased estimates of *F*_ST_ supported by the law of large numbers. Genomic data highlight the usefulness of the bias-corrected moment estimators of *F*_ST_.

## ACKNOWLEDGMENTS

We express our appreciation to editors Andrew Kern, Jeffrey Ross-Ibarra, Mark Beaumont, and Nicholas Barton for reviewing this manuscript and providing constructive comments. We also thank the reviewers for providing essential comments that significantly improved the earlier versions of the manuscript. This study was supported by Japan Society for the Promotion of Science Grants-in-Aid for Scientific Research KAKENHI nos. 16H02788 and 19H04070 to HK, and 18K0578116 to SK.

