## Supplemental Figures and Note for "Understanding population structure in an evolutionary context: population-specific *F*_ST_ and pairwise *F*_ST_"

This PDF file includes two sets of supplemental materials.

A. Supplemental Figures S1–S10
B. Supplemental Note

Supplemental Tables 1-7 are provided in an Excel file.

The R codes for our representation method exemplified by the human data and simulations of population colonization used in this study are available in the Supplemental Material.

### A. Supplemental Figures S1–S10

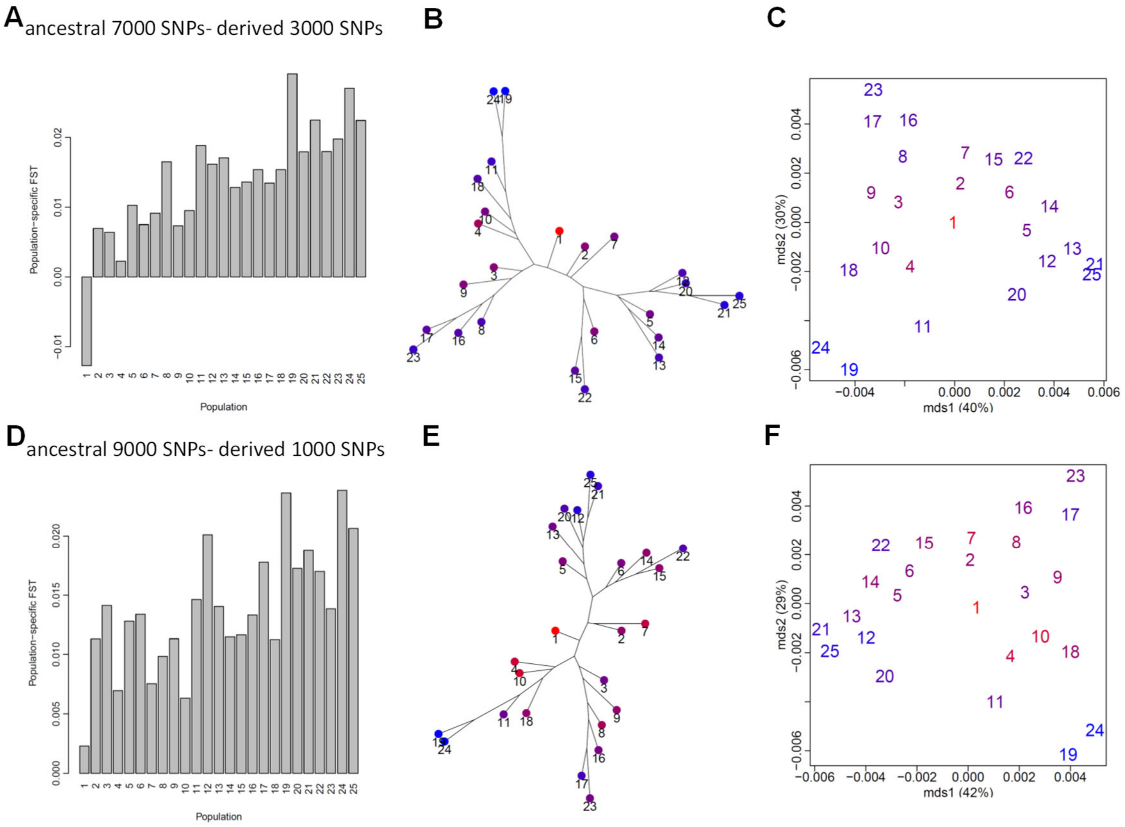


**Figure S1** Population colonization simulations of eight-directional grid colonization (Figure 1E). (Upper) 7,000 ancestral SNPs and 3,000 newly derived SNPs were selected. (Bottom) 9,000 ancestral SNPs and 1,000 newly derived SNPs were selected. (A, D) Population-specific $F_{ST}$ values. (B, E) Unrooted neighbor -joining (NJ) tree based on pairwise $F_{ST}$ overlaid with population-specific $F_{ST}$ values. (C, F) Multi-dimensional scaling (MDS) of pairwise $F_{ST}$. The color of each population indicates the magnitude of population-specific $F_{ST}$ values between red (for the smallest $F_{ST}$) and blue (for the largest $F_{ST}$).

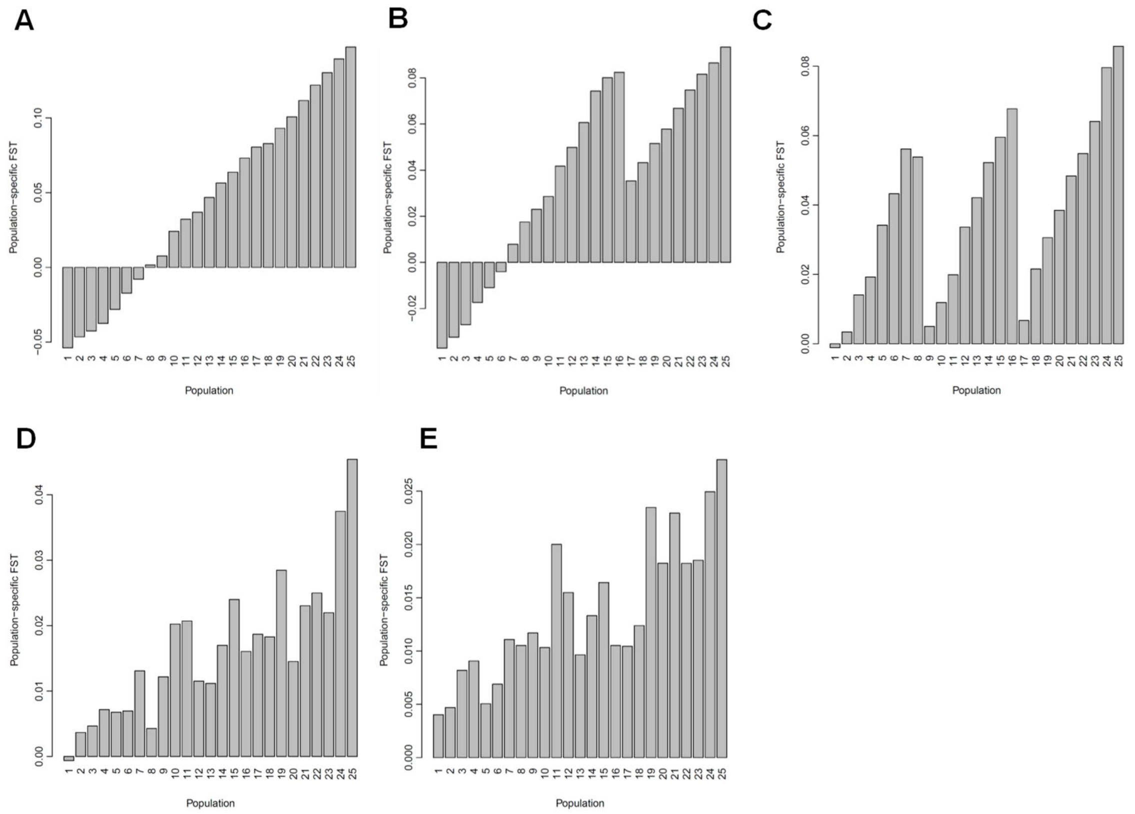


**Figure S2** Population-specific $F_{ST}$ values for 25 simulated populations are presented for each model in Figure 1 (A–E).

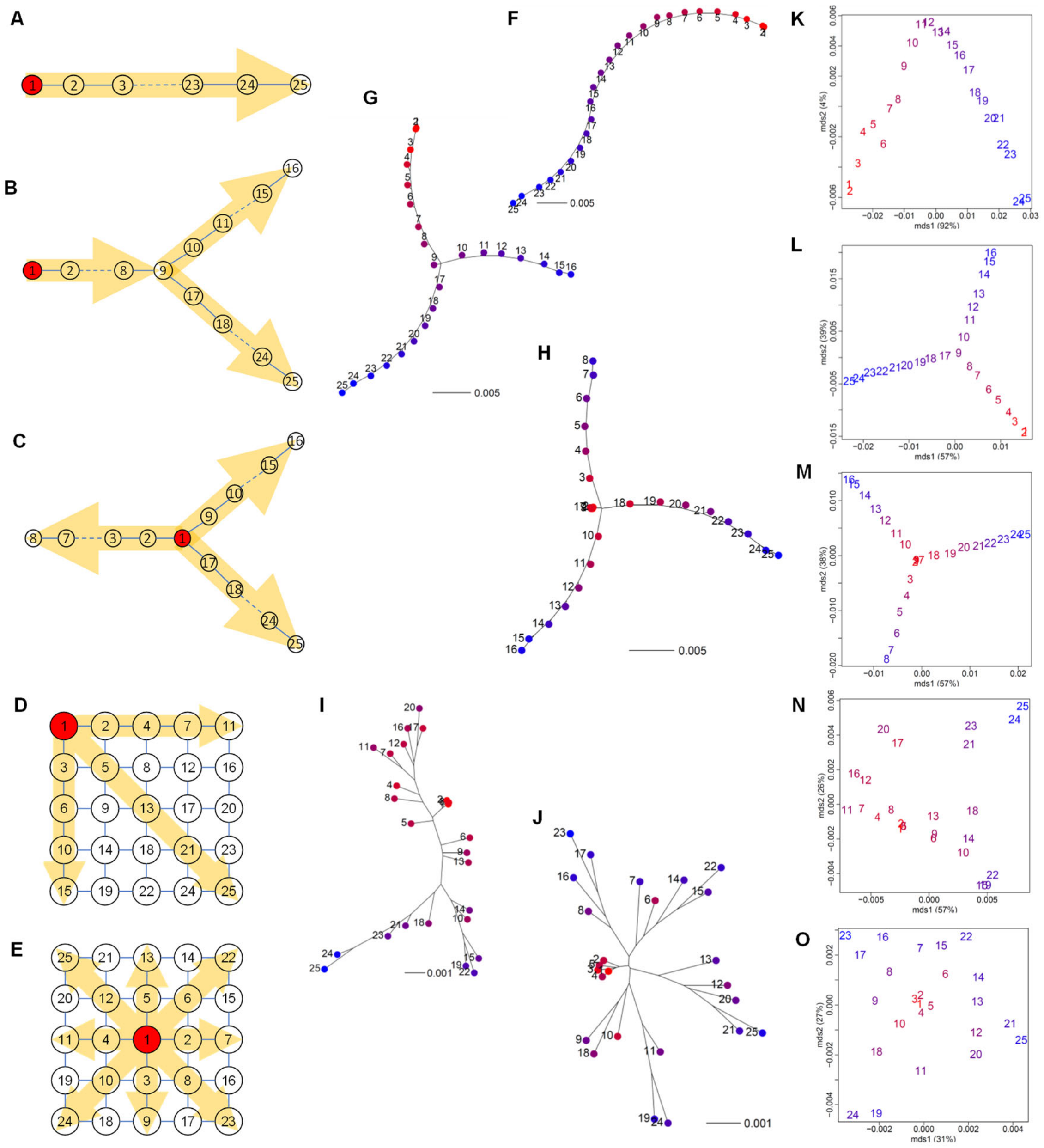

**Figure S3** Results from population colonization simulations. Schematic diagrams of the models: (A) one, (B) two, (C) three-directional colonization, (D) three-directional grid colonization from an edge, and (E) eight-directional grid colonization from the center. Population 1 in red is ancestral, and the yellow arrows indicate the direction of colonization. Lines show opportunities for migration. The effective population size of the ancestral population was 10 times greater ($N_e = 10^5$) than that of the newly derived population ($N_e = 10^4$) after one generation, and each population exchanged 1% of $N_e$ genes with adjacent population(s) in every generation, as indicated by the arrows (see the text). Neighbor-joining (NJ) unrooted trees (F–J) and multi-dimensional scaling (MDS) plots (K–O) based on the pairwise $F_{\mathrm{ST}}$ distance matrix overlaid with population-specific $F_{\mathrm{ST}}$ values for each model. The color of each population indicates the magnitude of population-specific $F_{\mathrm{ST}}$ values between red (for the smallest $F_{\mathrm{ST}}$) and blue (for the largest $F_{\mathrm{ST}}$).

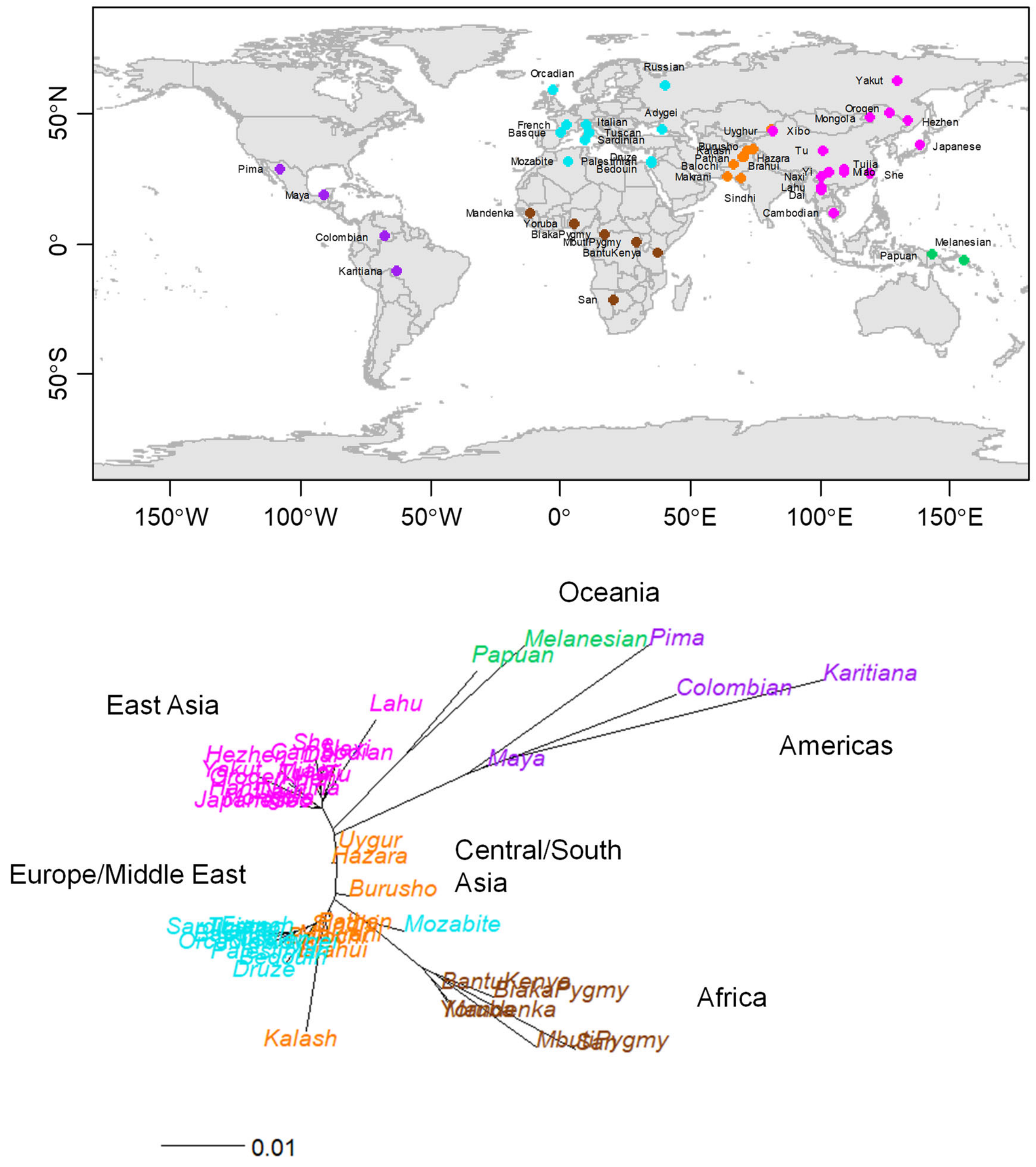


**Figure S4** Population structure of human populations. Sampling locations for 51 populations (upper). Data from Cann et al. (2002). Neighbor-joining (NJ) unrooted tree based on the pairwise $F_{ST}$ distance matrix (lower). Data from Rosenberg *et al.* (2002).

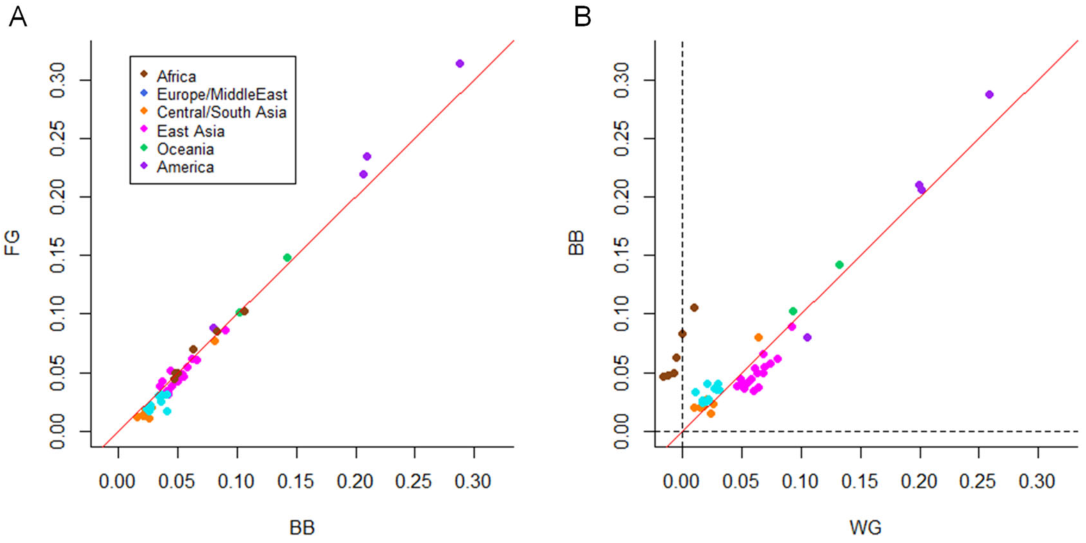


**Figure S5** Relationships between different population-specific $F_{ST}$ estimators for 51 human populations. (A) BB (Beaumont and Balding, 2004) vs. FG (Foll and Gaggiotti 2006) ($R^2 = 0.99, p = 2.2 \times 10^{-16}$). (B) WG (Weir and Goudet 2017) vs. BB ($R^2 = 0.87, p = 2.2 \times 10^{-16}$). Data from Rosenberg *et al.* (2002).

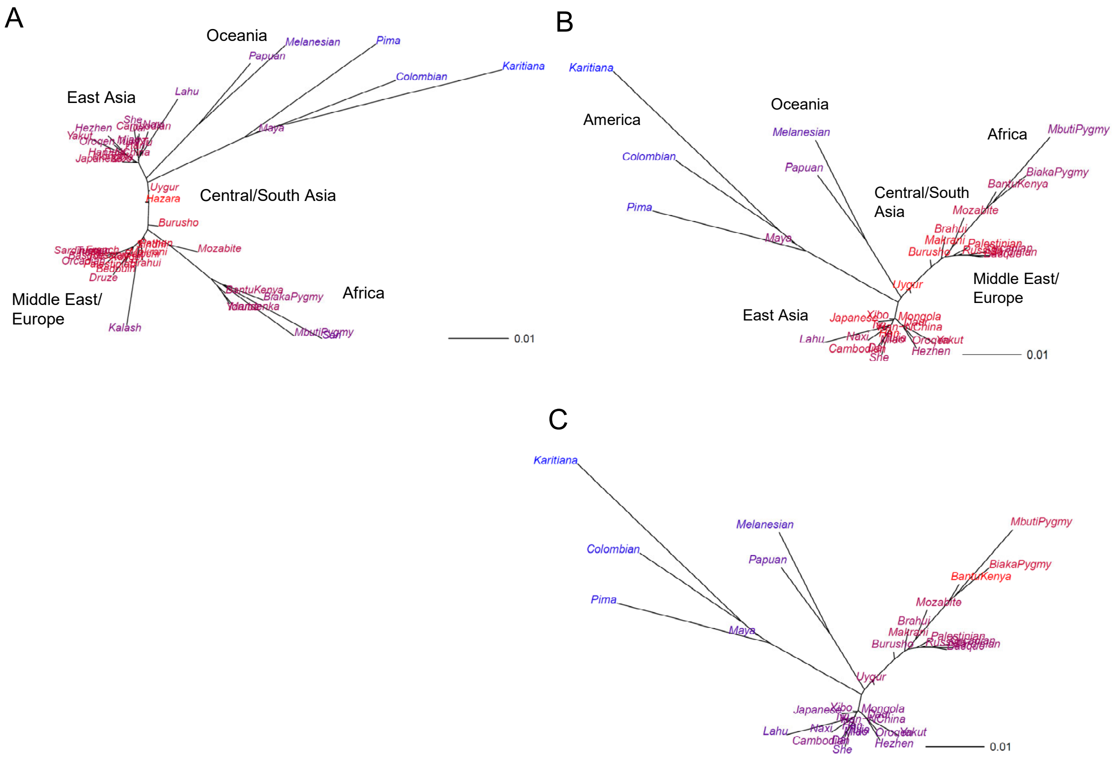


**Figure S6** Population structure of humans based on Bayesian (Beaumont and Balding, 2004) and moment population-specific $F_{ST}$ (Weir and Goudet 2017) estimators. Neighbor-joining (NJ) unrooted trees of the pairwise $F_{ST}$ distance matrix obtained from the Bayesian population-specific $F_{ST}$ estimator using (A) 51 samples and (B) 37 subsamples. (C) WG population-specific moment $F_{ST}$ using 37 subsamples. The numbers of sampling locations of the subsamples were as follows: 3 from Africa, 6 from the Middle East/Europe, 9 from Central/South Asia, 18 from East Asia, 2 from Oceania, and 4 from America. The color of each population indicates the magnitude of population-specific $F_{ST}$ values between red (for the smallest $F_{ST}$) and blue (for the largest $F_{ST}$).

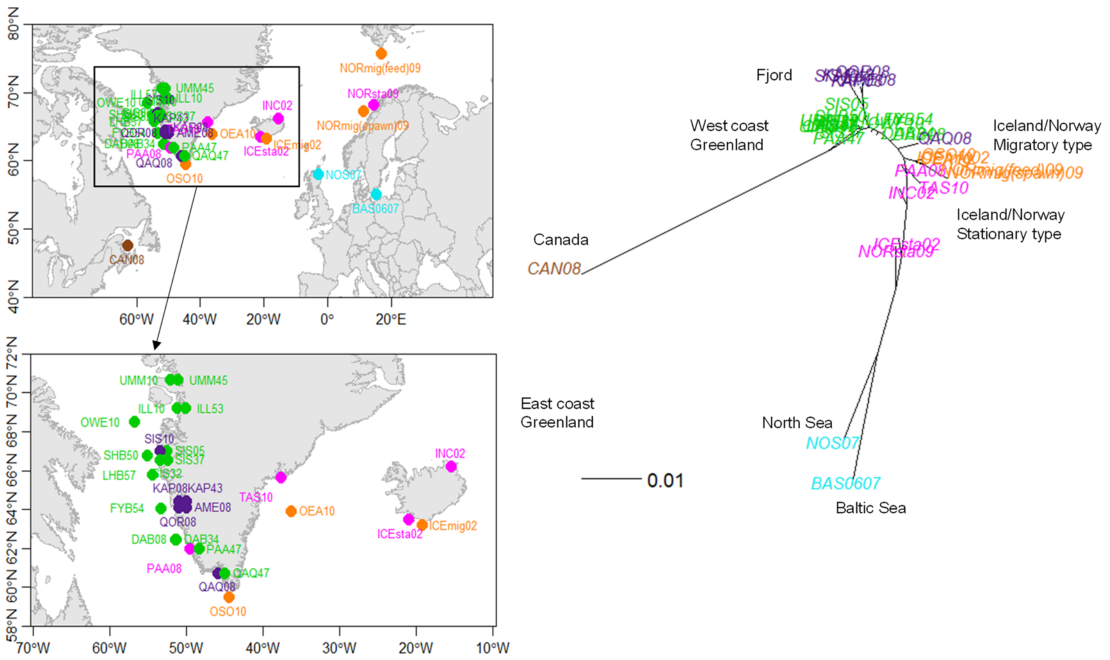


**Figure S7** Population structure of Atlantic cod populations. Sampling locations of 34 wild Atlantic cod populations (left). NJ unrooted tree based on the pairwise $F_{ST}$ distance matrix (right). Data from Therkildsen *et al.* (2013) and Hemmer-Hansen (2013).

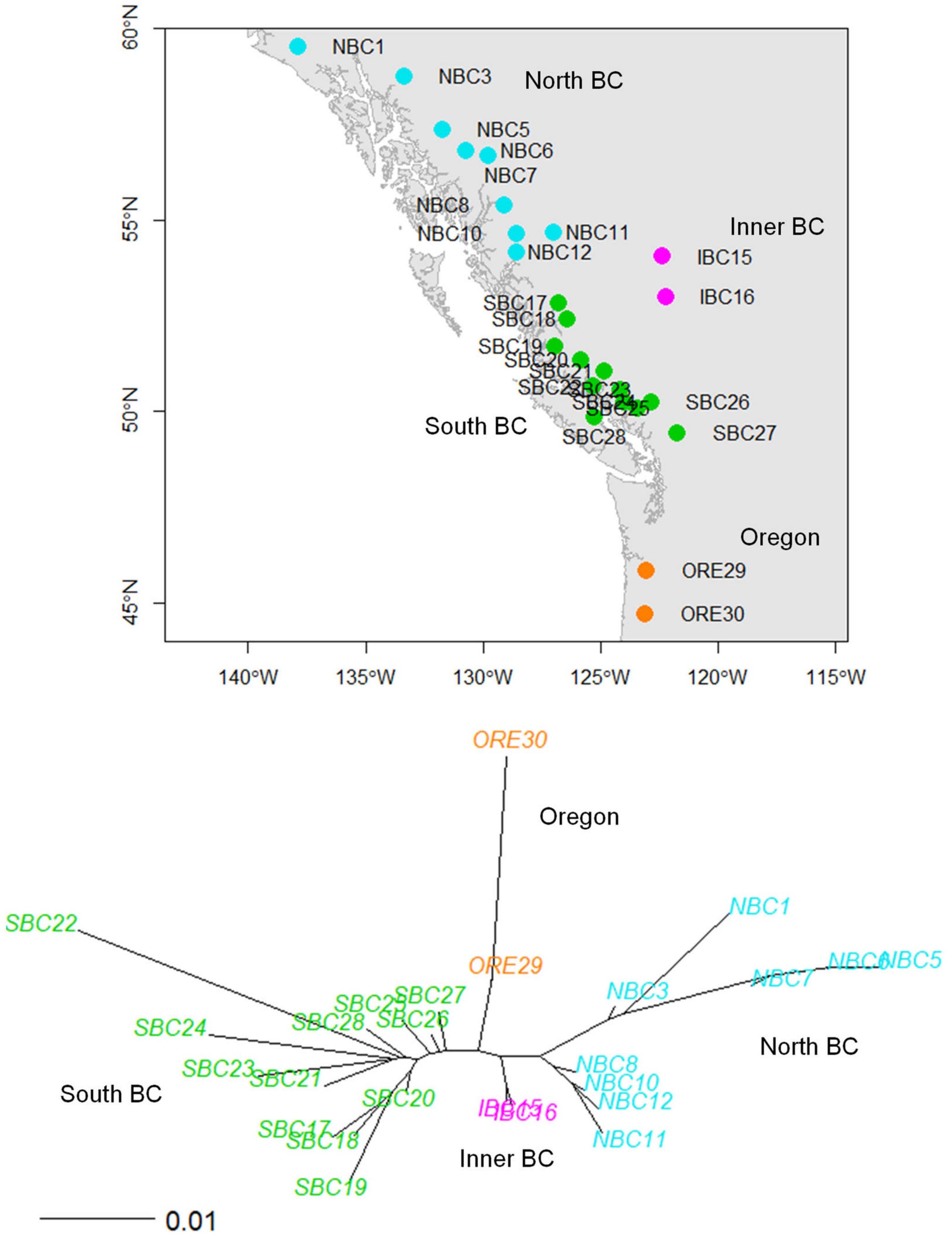


**Figure S8** Population structure of wild poplar populations. Sampling locations of 25 wild poplar populations (upper). Neighbor-joining (NJ) unrooted tree based on the pairwise $F_{ST}$ distance matrix (lower). Data from McKown *et al.* (2014b).

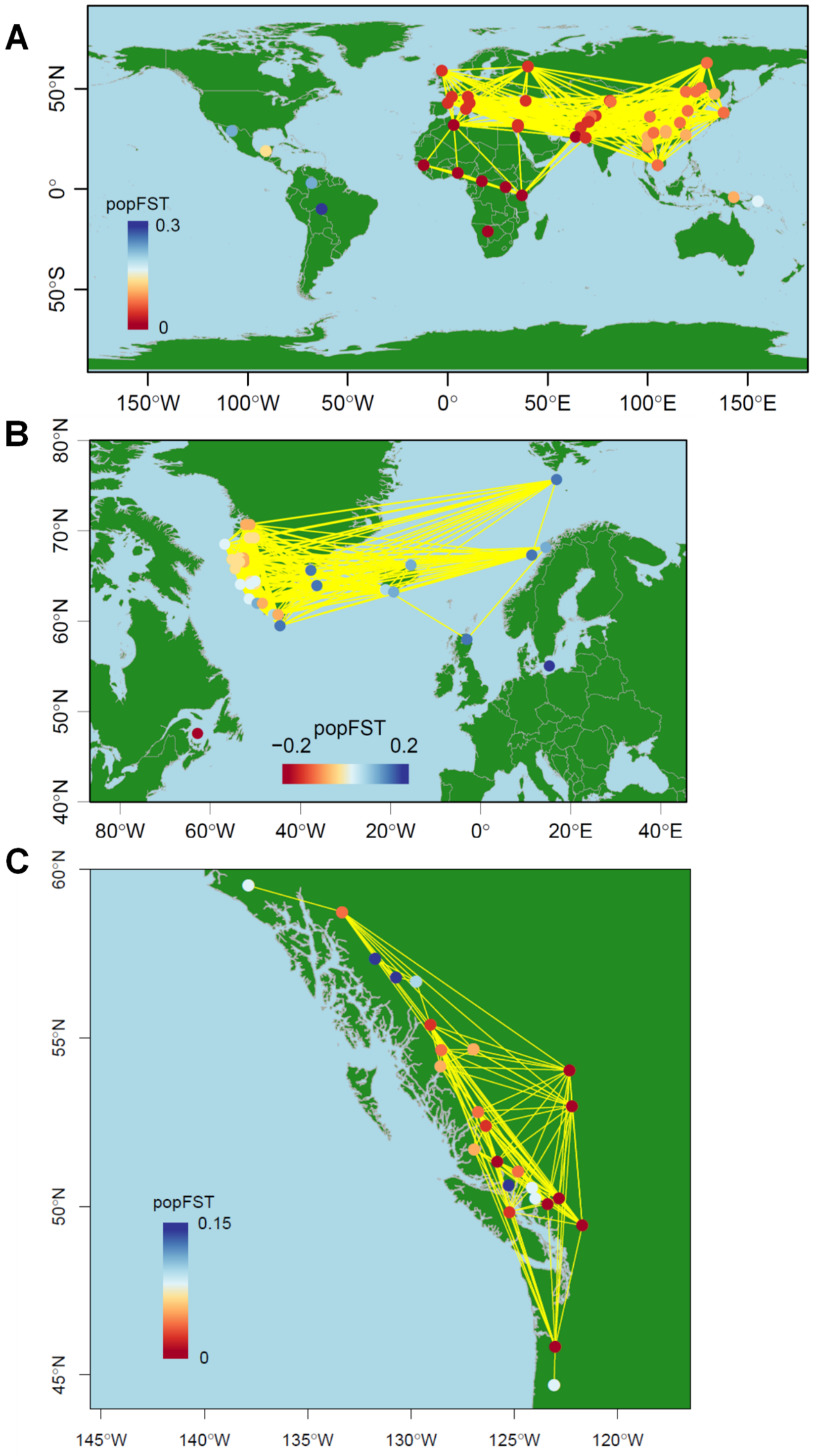


**Figure S9** Map showing population connectivity with the magnitude of population-specific $F_{ST}$ values using a diverging color palette (see the text). (A) Human. (B) Atlantic cod. (C) Wild poplar. Populations connected by yellow lines are those with pairwise $F_{ST} < 0.02$.

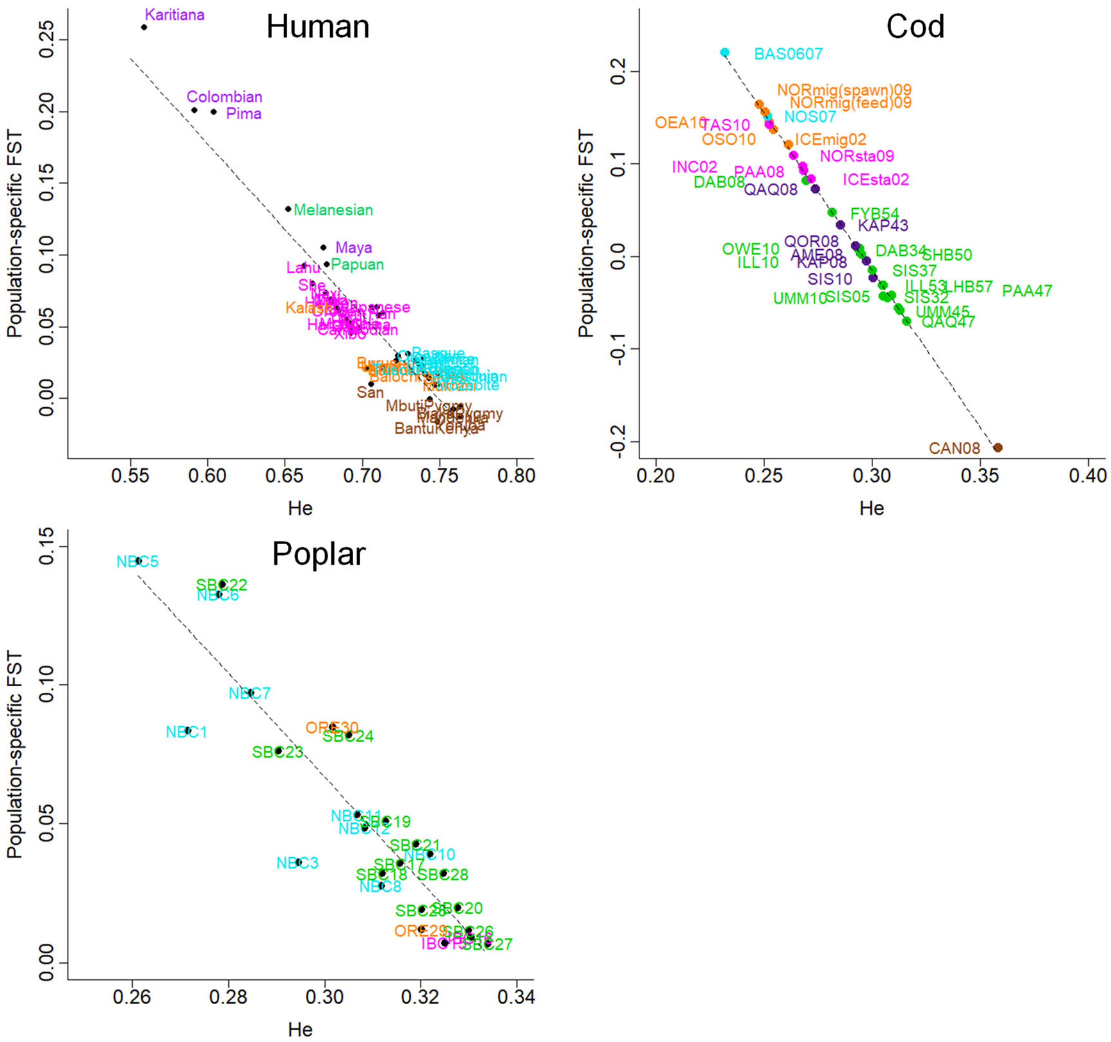


**Figure S10** *He* vs. Weir and Goudet's population-specific $F_{ST}$ values. Human: $y = -0.1895x + 0.8908$ ($R^2 = 0.91, F = 501.8$ (1 ,49DF), $P < 2.2 \times 10^{-16}$ ). Atlantic cod: $y = -3.397x + 1.004$ ($R^2 = 0.998, F = 20250$ (1 ,32DF), $P < 2.2 \times 10^{-16}$). Wild poplar: $y = -1.872x + 0.629$ ($R^2 = 0.82, F = 108$ (1 ,23DF), $P < 3.5 \times 10^{-10}$).

#### B. Supplemental Note

1. $G_{ST}$ and Nei and Chesser's $F_{ST}$ estimator
2. Weir and Goudet's population-specific $F_{ST}$ estimator
3. $F_{ST}$ estimators used in this study

We summarized the $F_{ST}$ estimators used in our study for readers' convenience, using notations consistent with those of Weir and Hill (2002): *i* for populations ($i = 1, \dots, r$), *u* for alleles ($u = 1, \dots, m$), and *l* for loci ($l = 1, \dots, L$) (see original papers that developed the $F_{ST}$ estimators for detailed information).

##### 1. $G_{ST}$ and Nei and Chesser's $F_{ST}$ estimator

Nei (1973) proposed the $G_{ST}$ measure to explicitly formulate Wright's *F*-statistics using genetic diversity while allowing multiple alleles. $G_{ST}$ is equal to $F_{ST}$ for diploid random mating populations (Excoffier 2007).

$G_{ST}$ is defined as the ratio of between-population heterozygosity to total heterozygosity; that is,

$$G_{\mathrm{ST}} = \frac{H_T - H_S}{H_T}$$

where $H_T$ is total heterozygosity and $H_S$ is within-population heterozygosity. Here, all populations are assumed in Hardy-Weinberg Equilibrium (Nei, 1973). At a single locus, $H_{\mathrm{T}}$ (total heterozygosity) and $H_{\mathrm{S}}$ (within-population heterozygosity) are

$$H_S = 1 - \frac{1}{r}\sum_{i=1}^{r}\sum_{u=1}^{m} p_{iu}^2$$

$$H_T = 1 - \sum_{u=1}^{m}\left(\frac{1}{r}\sum_{i=1}^{r} p_{iu}\right)^2$$

$F_{ST}$ is the ratio of between to total variance. The estimator is therefore a ratio estimator and consequently biased. To correct the bias, Nei and Chesser (1983) derived the unbiased moment estimators of $H_S$ and $H_T$ in diploid populations. This estimator (NC83) assumes that samples ($n_i$ individuals) are randomly chosen from a set of fixed populations ($i = 1, \dots, r$). The gene diversities in population *i* are written based on the homozygote genotype frequency $P_{iuu}$ and allele frequency $p_{iu}$, which are given as

$$H_{0i} = 1 - \sum_{u=1}^{m} P_{iuu}$$

$$H_{Si} = 1 - \sum_{u=1}^{m} p_{iu}^2$$

Because observed genotype frequencies are unbiased, $H_{0i}$ is unbiasedly estimated by

$$\widehat{H}_{0i} = 1 - \sum_{u=1}^{m} \tilde{P}_{iuu}$$

where $\tilde{P}_{iuu}$ denotes observed homozygote genotype frequencies. To estimate $H_{Si}$, Nei and Chesser (1983) used the expectation of $\tilde{p}_{iu}^2$, because $\tilde{p}_{iu}^2$ is biased, as

$$E[\tilde{p}_{iu}^2] = p_{iu}^2 + \frac{P_{iuu}}{n_i} + \sum_{u \neq u}^{m} \frac{P_{iuu'}}{4n_i} - \frac{p_{iu}^2}{n_i}$$

Those authors then calculated the expectation of observed gene diversity in population *i*: because $p_{iu} = P_{uu} + \frac{\sum_{u \neq u}^{m} P_{uu'}}{2}$,

$$1 - E\left[\sum_{u=1}^{m} \tilde{p}_{iu}^2\right] = 1 - \sum_{u=1}^{m} p_{iu}^2 - \frac{1}{n_i}\sum_{u=1}^{m} P_{iuu} - \frac{1}{2n_i}\sum_{u=1}^{m}(p_{iu} - P_{uu}) + \frac{1}{n_i}\sum_{u=1}^{m} p_{iu}^2$$

$$= 1 - \sum_{u=1}^{m} p_{iu}^2 - \frac{1}{n_i}\sum_{u=1}^{m} P_{iuu} - \frac{1}{2n_i} + \frac{1}{2n_i}\sum_{u=1}^{m} P_{uu} + \frac{1}{n_i}\sum_{u=1}^{m} p_{iu}^2$$

$$= H_{Si}\left(1 - \frac{1}{n_i}\right) + \frac{H_{0i}}{2n_i} \qquad \text{(NC83 Eq. 6)}$$

Using the method of moments, they obtained the unbiased estimator of gene diversity in population $i$ $(H_{Si})$:

$$\hat{H}_{Si} = \frac{n_i}{n_i - 1}\left(1 - \sum_{u=1}^{m} \tilde{p}_{iu}^2 - \frac{\hat{H}_{0i}}{2n_i}\right) \qquad \text{(NC83 Eq. 7)}$$

Because observed genotype frequencies are unbiased, $H_{0i}$ was unbiasedly estimated by $\hat{H}_{0i} = 1 - \sum_{u=1}^{m} \tilde{P}_{iuu}$. Thus, they unintentionally derived population-specific gene diversity.

Under the assumption that all populations are in Hardy–Weinberg equilibrium, $\hat{H}_{Si} = \hat{H}_{0i}$, we obtain

$$\hat{H}_{Si} = \frac{2n_i}{2n_i - 1}\left(1 - \sum_{u=1}^{m} \tilde{p}_{iu}^2\right) \qquad \text{(Eq. S1)}$$

The expectation of the average of observed gene diversity over all populations was similarly derived as

$$1 - E\left[\sum_{u=1}^{m} \overline{\tilde{p}_u^2}\right] = \frac{1}{r}\left[\sum_{i=1}^{r}\left\{H_{Si}\left(1 - \frac{1}{n_i}\right) + \frac{H_{0i}}{2n_i}\right\}\right]$$

$$= H_S\left(1 - \frac{1}{\tilde{n}}\right) + \frac{H_0}{2\tilde{n}} \qquad \text{(NC83 Eq. 8)}$$

where $\overline{\tilde{p}_u^2} = \frac{1}{r}\sum_{i=1}^{r} \tilde{p}_{iu}^2$, and $\tilde{n}$ is the harmonic mean of $n_i$, namely, $\tilde{n} = \frac{r}{\sum_{i=1}^{r}\frac{1}{n_i}}$ . The unbiased moment estimator of gene diversity over all populations was derived as

$$\hat{H}_S = \frac{\tilde{n}}{\tilde{n} - 1}\left(1 - \sum_{u=1}^{m} \overline{\tilde{p}_u^2} - \frac{\hat{H}_0}{2\tilde{n}}\right) \qquad \text{(NC83 Eq. 9)}$$

Here, $\hat{H}_0$ is the unbiased estimator of gene diversity ($H_o = 1 - \sum_{u=1}^{m} P_{uu}$) based on the homozygote genotype frequencies over all populations:

$$\hat{H}_0 = 1 - \sum_{u=1}^{m} \bar{\tilde{P}}_{uu} \qquad \text{(NC83 Eq. 5)}$$

where $\bar{\tilde{P}}_{uu} = \frac{1}{r}\sum_{i=1}^{r} \tilde{P}_{uu}$. To obtain the unbiased estimator of $H_{\mathrm{T}}$, they derived the expectation of observed mean gene diversity over all populations:

$$1 - E\left[\sum_{u=1}^{m} \bar{\tilde{p}}_u^2\right] = H_T - \frac{H_S}{\tilde{n}r} + \frac{H_0}{2\tilde{n}r} \qquad \text{(NC83 Eq. 10)}$$

where $\bar{\tilde{p}}_u = \frac{1}{r}\sum_{i=1}^{r} \tilde{p}_{iu}$. From this equation, they obtained the unbiased moment estimator of $H_T$ as

$$\hat{H}_T = 1 - \sum_{u=1}^{m} \bar{\tilde{p}}_u^2 + \frac{\hat{H}_S}{\tilde{n}r} - \frac{\hat{H}_0}{2\tilde{n}r} \qquad \text{(NC83 Eq. 11)}$$

Under the assumption that all populations are in Hardy–Weinberg equilibrium, $H_0 = H_S$. In this situation, the estimators are a function of allele frequencies only at a single locus:

$$\hat{H}_S = \frac{2\tilde{n}}{2\tilde{n}-1}\left(1 - \sum_{u=1}^{m} \overline{\tilde{p}_u^2}\right) \qquad \text{(NC83 Eq. 15)}$$

$$\hat{H}_T = 1 - \sum_{u=1}^{m} \bar{\tilde{p}}_u^2 + \frac{\hat{H}_S}{2\tilde{n}r} \qquad \text{(NC83 Eq. 16)}$$

#### 2. Weir and Goudet's population-specific $F_{\mathrm{ST}}$ estimator

Weir and Goudet (2017) derived the bias-corrected moment estimator of population-specific $F_{\mathrm{ST}}$ (WG) for random mating populations, when only allele frequencies are used, as

$$\mathrm{ps}\hat{F}_{\mathrm{ST}}^{i} = \hat{\beta}_{\mathrm{WT}}^{i} = \frac{\tilde{M}_w^i - \tilde{M}^B}{1 - \tilde{M}^B} \qquad \text{(WG Table 3)}$$

$\tilde{M}_W^i$ is the unbiased within-population matching of two distinct alleles of population *i:*

$$\tilde{M}_W^i = \frac{2n_i}{2n_i - 1}\sum_{u=1}^{m} \tilde{p}_{iu}^2 - \frac{1}{2n_i - 1} \qquad \text{(WG Table 2)}$$

where $n_i$ is the sample size (number of individuals) taken from population $i$, and $\tilde{p}_{iu}$ is the observed frequency of allele *u*. We note that $1 - \tilde{M}_W^i$ equals $\hat{H}_{Si}$ (Equation S1):

$$1 - \tilde{M}_W^i = \frac{2n_i}{2n_i - 1}\left(1 - \sum\nolimits_{u=1}^{m} \tilde{p}_{iu}^2\right) = \hat{H}_{Si} \qquad \text{(Eq. S2)}$$

$\tilde{M}^B$ is the between-population-pair matching average over pairs of populations $i, i'$:

$$\tilde{M}^B = \frac{1}{r(r-1)}\sum_{i=1}^{r}\sum_{i'=1(i'\neq i)}^{r} \tilde{M}_B^{ii'} \qquad \text{(WG Table 2)}$$

$\tilde{M}_B^{ii'}$ is the matching of one allele in $j, j'$ individuals taken from each of populations $i, i'$:

$$\tilde{M}_B^{ii'} = \frac{1}{n_i n_{i'}}\sum_{j=1}^{n_i}\sum_{j'=1}^{n_{i'}} \tilde{M}_{jj'}^{ii'} = \sum_{u=1}^{m} \tilde{p}_{iu}\tilde{p}_{i'u} \qquad \text{(WG Table 2)}$$

#### 3. $F_{\mathrm{ST}}$ estimators applied in this study

To estimate genome-wide pairwise $F_{\mathrm{ST}}$, we used the NC83 bias-corrected $G_{\mathrm{ST}}$ moment estimator (Nei and Chesser 1983) for overall loci ($l = 1, \ldots, L$) (genome-wide pairwise $F_{\mathrm{ST}}$):

$$\mathrm{pw}\hat{F}_{\mathrm{ST}}^{ij} = \frac{\sum_{l=1}^{L}\left(\hat{H}_{T,l}^{ij} - \hat{H}_{S,l}^{ij}\right)}{\sum_{l=1}^{L} \hat{H}_{T,l}^{ij}} \qquad \text{(Eq. S3)}$$

In our analyses, we extended the WG population-specific $F_{\mathrm{ST}}$ estimator (Weir and

Goudet 2017) to overall loci (genome-wide population-specific $F_{\mathrm{ST}}$) (Buckleton *et al.* 2016):

$$\mathrm{ps}\hat{F}_{\mathrm{ST}}^{i} = \frac{\sum_{l=1}^{L}(\widetilde{M}_{W,l}^{i} - \widetilde{M}_{l}^{B})}{\sum_{l=1}^{L}(1 - \widetilde{M}_{l}^{B})} \qquad (\mathrm{Eq.\,S4})$$

These are called the "ratio of averages" $F_{\mathrm{ST}}$ estimator (Weir and Cockerham 1984; Weir and Hill 2002; Bhatia et al. 2013).

The variance–covariance matrix of the population-specific $F_{\mathrm{ST}}$ estimator can be written as $\mathrm{ps}\hat{F}_{\mathrm{ST}}^{i} = \frac{\bar{y}^{i}}{\bar{x}}$. Using the Taylor series expansion for the first term, we inferred the asymptotic variance as

$$V\left[\mathrm{ps}\hat{F}_{\mathrm{ST}}^{i}\right] \simeq \frac{\bar{y}^{2}}{\bar{x}^{2}}\left\{\frac{V[\bar{x}]}{\bar{x}^{2}} + \frac{V\left[\bar{y}^{i}\right]}{\bar{y}^{i^{2}}} - \frac{2Cov\left[\bar{x},\bar{y}^{i}\right]}{\bar{x}\bar{y}^{i}}\right\} \qquad (\mathrm{Eq.\,S5})$$

Similarly, the asymptotic covariances between population-specific $F_{\mathrm{ST}}$ values of $i, j$ populations were obtained by

$$Cov\left[\mathrm{ps}\hat{F}_{\mathrm{ST}}^{i}, \mathrm{ps}\hat{F}_{\mathrm{ST}}^{j}\right] \simeq \frac{\bar{y}^{i}\bar{y}^{j}}{\bar{x}^{4}}V[\bar{x}] - \frac{\bar{y}^{i}}{\bar{x}^{3}}Cov\left[\bar{x},\bar{y}^{j}\right] - \frac{\bar{y}^{j}}{\bar{x}^{3}}Cov\left[\bar{x},\bar{y}^{i}\right] \qquad (\mathrm{Eq.\,S6})$$

where the variance and covariance components were calculated by

$V[\bar{x}] = \frac{1}{L(L-1)}\sum_{l=1}^{L}(x_l - \bar{x})^2$ , $V\left[\bar{y}^{i}\right] = \frac{1}{L(L-1)}\sum_{l=1}^{L}\left(y_l^{i} - \bar{y}^{i}\right)^2$,

$Cov\left[\bar{x},\bar{y}^{i}\right] = \frac{1}{L(L-1)}\sum_{l=1}^{L}(x_l - \bar{x})\left(y_l^{i} - \bar{y}^{i}\right)$ ,

$Cov\left[\bar{x},\bar{y}^{j}\right] = \frac{1}{L(L-1)}\sum_{l=1}^{L}(x_l - \bar{x})\left(y_l^{j} - \bar{y}^{j}\right)$

We also applied empirical (Beaumont and Balding, 2004) and full Bayesian (Foll and Gaggiotti, 2006) population-specific $F_{\mathrm{ST}}$ estimators. Beaumont and Balding (2004) maximized the Dirichlet-multinomial marginal likelihood in their Equation 1 and estimated $\theta_{li}$:

$$L_{li}\left(\theta_{li}\middle|n_{li1},\ldots,n_{lim_l}\right) = \frac{\Gamma(\theta_{li})}{\Gamma(N_{li} + \theta_{li})}\prod_{u=1}^{m_l}\frac{\Gamma(n_{liu} + \theta_{li}\bar{p}_{lu})}{\Gamma(\theta_{li}\bar{p}_{lu})} \qquad (\mathrm{Eq.\,S7})$$

Here, $\theta_{li}$ is the scale parameter of the Dirichlet prior distribution for locus *l* and population $i$, $\tilde{p}_{lu}$ is the observed frequency of allele *u* ($u = 1,\ldots,m$) at locus *l*, $n_{liu}$ is the observed allele count in population $i$, and $N_{li}$ is the total number of alleles. Importantly, $\bar{p}_{lu}$ is the mean allele frequency over all populations, whereas $\theta_{li}\bar{p}_{lu} = \alpha_{liu}$, where $\theta_{li} = \sum_{u=1}^{m_l}\alpha_{liu}$. The parametrization reduces the number of parameters to be estimated. Based on a Dirichlet (multi-allelic) and/or a beta (bi-allelic) scale parameter, population-specific $F_{\mathrm{ST}}$ values were estimated for each locus using the following function of $\hat{\theta}_{li}$ (Beaumont and Balding, 2004):

$$\mathrm{ps}\tilde{F}_{\mathrm{ST},l}^{i} = \frac{1}{\hat{\theta}_{li} + 1} \qquad (\mathrm{Eq.\,S8})$$
